# Identification of multiple kinetic populations of DNA-binding proteins in live cells

**DOI:** 10.1101/509620

**Authors:** Han N. Ho, Daniel Zalami, Jürgen Köhler, Antoine M. van Oijen, Harshad Ghodke

## Abstract

Understanding how multi-protein complexes function in cells requires detailed quantitative understanding of their association and dissociation kinetics. Analysis of the heterogeneity of binding lifetimes enables interrogation of the various intermediate states formed during the reaction. Single-molecule fluorescence imaging permits the measurement of reaction kinetics inside living organisms with minimal perturbation. However, poor photo-physical properties of fluorescent probes limit the dynamic range and accuracy of measurements of off rates in live cells. Time-lapse single-molecule fluorescence imaging can partially overcome the limits of photobleaching, however, limitations of this technique remain uncharacterized. Here, we present a structured analysis of which timescales are most accessible using the time-lapse imaging approach and explore uncertainties in determining kinetic sub-populations. We demonstrate the effect of shot noise on the precision of the measurements, as well as the resolution and dynamic range limits that are inherent to the method. Our work provides a convenient implementation to determine theoretical errors from measurements and to support interpretation of experimental data.

**STATEMENT OF SIGNIFICANCE:** Measuring lifetimes of interactions between DNA-binding proteins and their substrates is important for understanding how they function in cells. In principle, time-lapse imaging of fluorescently-tagged proteins using single-molecule methods can be used to identify multiple sub-populations of DNA-binding proteins and determine binding lifetimes lasting for several tens of minutes. Despite this potential, currently available guidelines for the selection of binding models are unreliable, and the practical implementation of this approach is limited. Here, using experimental and simulated data we identify the minimum size of the dataset required to resolve multiple populations reliably and measure binding lifetimes with desired accuracy. This work serves to provide a guide to data collection, and measurement of DNA-binding lifetimes from single-molecule time-lapse imaging data.

## INTRODUCTION

Understanding fundamental processes of life requires characterization of the kinetics of interactions between biological molecules. At single-molecule levels, these systems often exhibit kinetic heterogeneity that is inherent to the presence of multiple intermediate states (1–17). Advances in single-molecule imaging have enabled the detection and characterization of heterogeneous sub-populations in reactions conducted *in vitro* as well as, *in vivo.* Ultimately, these investigations enable the construction of detailed molecular mechanisms to explain how various biomolecular interactions proceed.

Compared to *in vitro* studies, live-cell investigations offer the key advantage of studying biochemical reactions at physiological conditions that can be difficult to reconstitute. Singlemolecule live-cell imaging commonly relies on fluorescent proteins that are genetically fused to the protein of interest (Fig. 1A) (18–22). Tracking the fluorescence signal of thousands of molecules, one molecule at a time, enables the building of physical models, from which physical parameters such as diffusion constants and detachment rates from DNA can be determined. Where detachment rates are concerned, the trajectory lengths of thousands of molecules are aligned to obtain a cumulative residence time distribution (CRTD). At the single-molecule level, the dissociation of a protein from its substrate is a stochastic process. This phenomenon can be adequately described as a two-state kinetic model with the interconversion of populations being modelled as a Poisson process. The resulting CRTD can be fit to exponential functions to obtain decay rates. In the case of a fluorescently tagged protein where loss of fluorescence is attributable to either dissociation, or photobleaching of the chromophore, the decay rate represents a combination of dissociation rates and photobleaching rate (Fig. 1B-C) (23).

**FIGURE 1.**
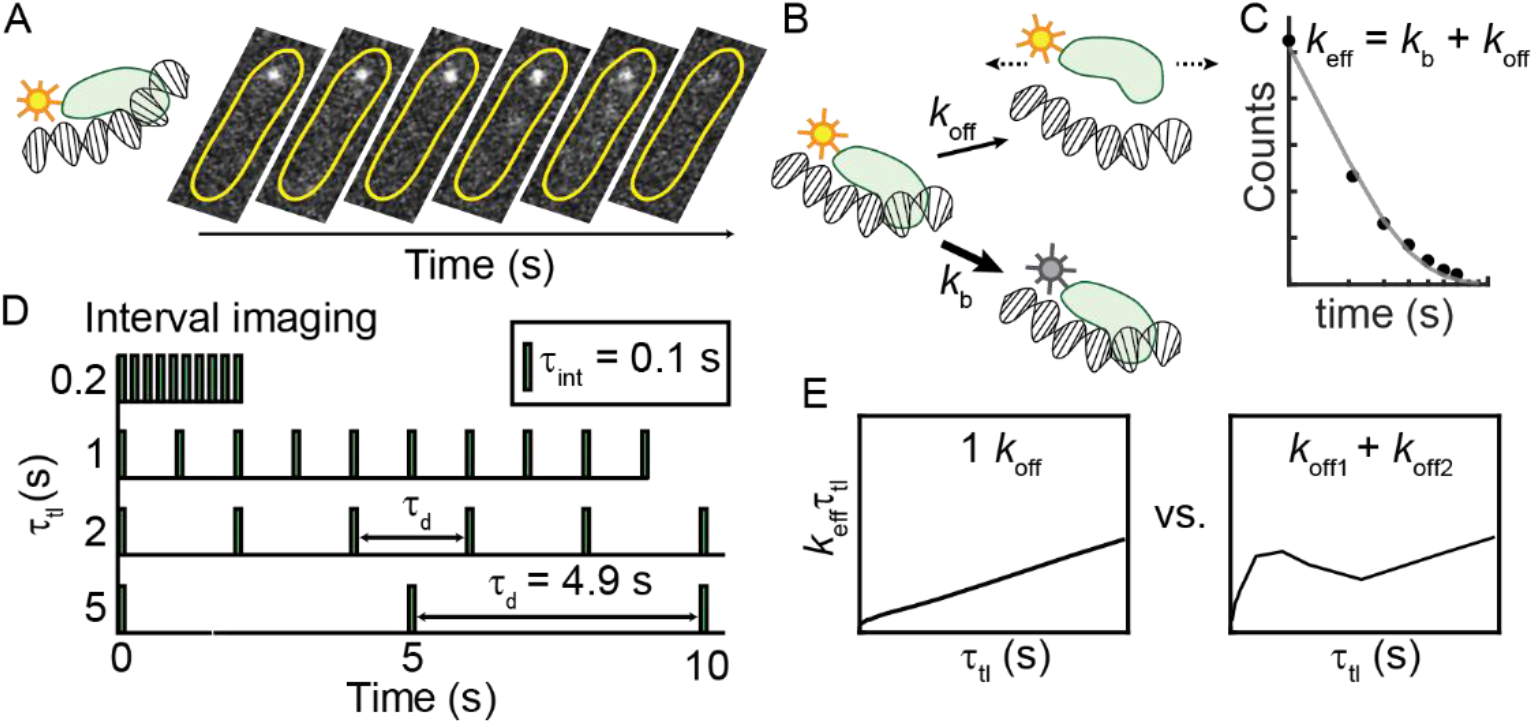
Experimental approach for characterizing kinetic heterogeneity of protein binding in live cells using single-molecule fluorescence imaging. (A) The protein of interest is tagged with a fluorescent protein. When the protein binds to DNA substrate, its fluorescence signal appears as a diffraction-limited focus that can be tracked in real time. Subsequent dissociation results in the disappearance of the focus and a redistribution of fluorescence signal throughout the cell. Yellow outlines illustrate the bacterial cell membrane. (B) The loss of fluorescence is attributable to either dissociation, or photobleaching of the chromophore. (C) Cumulative residence time distribution (CRTD) constructed from binding durations of thousands of molecules. Fitting the exponential function (Eq. 1) to CRTD yields an effective rate *k*_eff_, which is the sum of off rate (*k*_off_) of the protein of interest and photobleaching rate (*k*_b_) of the fluorescent probe (23). (D) To deconvolute *k*_b_ and *k*_off_, excitation and integration durations (τ_int_) can be spaced with various dark intervals (τ_d_). (E) Through exponential analyses, CRTDs obtained at various intervals result in *k*_eff_τ_tl_ plots which are indicators of kinetic heterogeneity (23). A single kinetic population yields a straight line whereas deviations from linear fits indicate the presence of a second kinetic sub-population. For a single kinetic population, the slope is the off rate and y-intercept is proportional to the photobleaching rate.

Photobleaching, a result of fluorescent proteins being damaged upon exposure to excitation sources, leads to the loss of fluorescence signal (24). Under excitation conditions that guarantee good signal-to-background ratios, fluorescent proteins can only stay ‘on’ for a few frames during continuous acquisitions. This limited visualization window reflects the ‘photon budget’ (25). Thus, when photobleaching occurs faster than the dissociation process, lifetime measurements are limited by the photobleaching rate. To overcome this problem and extend the observation time, the observation time window can be expanded by temporally spacing the photon budget using stroboscopic imaging (26). In this method, a dark interval (τ_d_) is inserted between integration time (τ_int_), effectively scaling the observation time with a factor of τ_tl_/τ_int_ (τ_tl_ = τ_int_ + τ_d_). Instead of using one dark interval, Gebhardt and co-workers (2013) developed an approach involving ‘time-lapse illumination with a fixed integration time, interspersed with dark periods of varying duration’ in which fluorescence acquisitions are collected at a series of time-lapse intervals (Fig. 1D) (23, 27). This method has also been variously referred to as ‘time-lapse imaging’ (28), ‘time-lapse illumination with different dark times’ (29), ‘time-lapse imaging at multiple timescales’ (30) and ‘stroboscopic single particle tracking PALM’ (31). For the purpose of brevity, and to distinguish from a time-lapse imaging mode with a single dark interval, we have adopted the term ‘interval imaging’ in our lab (32). Briefly, the approach works as follows: First, several movies (each with a unique dark interval) are collected while keeping the photon budget constant (in practice this is achieved by keeping the number of frames constant across all the movies). In cases where the copy number of the tagged protein is high and single-molecule imaging conditions may be difficult to attain, the cellular fluorescence is first photobleached such that only single-molecule fluorescence is observable. Subsequently, using particle tracking algorithms that enable measurements of lifetimes of bound molecules within a specified localization radius, a CRTD can be compiled. Fitting the CRTDs to effective rates (*k*_eff_), one can obtain the so-called *k*_eff_τ_tl_ plot which is linear for monoexponential distributions (Fig. 1E) (23). In this case, since the photobleaching rate is maintained constant across all conditions, it can be read off from the intercept on the Y-axis. A population of molecules dissociating with a finite and measurable off rate manifests as a straight line, where the slope reports on the off rate of the dissociation kinetics. A mixed population composed of species dissociating with multiple lifetimes manifests as a deviation from the linear fit (Fig. 1E) (23). Fitting the experimental data to a model describing mixed populations can then be used to extract the relative amplitudes and rates of the various populations. This power to deconvolute the photobleaching rate from multiple off rates has been successfully harnessed to dissect the kinetic heterogeneity of various DNA binding proteins including transcription factors and DNA replication and repair proteins in live cells (23, 27–33).

However, limitations arising from the practical implementation of this elegant method remain uncharacterized. In particular, we address the following questions: 1) What is the minimum number of observations needed to determine the binding lifetime of a species within a specified confidence? 2) For a given experimental setup, what is the dynamic range in binding lifetimes that can be detected? 3) How many populations can be resolved? and 4) What limits the ability to reliably resolve multiple populations? We consider four cases below to answer these questions. This study serves to provide a practical guide to realize the power as well as limitations of practical implementations of the interval imaging approach to measure intracellular binding kinetics of fluorescently tagged proteins.

## METHODS

### Rationale and model

For an introduction to the method, we direct the reader to seminal work by Gebhardt and coworkers who have developed and demonstrated the time-lapse imaging approach discussed here (23). Here, we first summarize the theoretical development to establish the context of the problem for this report. Consider a system containing *‘A’* number of fluorescently tagged DNA-bound proteins, wherein the proteins dissociate from DNA with a single off rate (*k*_off_). Upon exposure to excitation photon sources, the fluorescent proteins exhibit photobleaching with a rate kb, resulting in the loss of fluorescence signal. Additionally, dissociation contributes to the loss of fluorescent foci as protein molecules move out of the localization radius. Since dissociation and photobleaching are independent, and both are Poisson processes, the loss of observations as a function of time *t* can be described as:

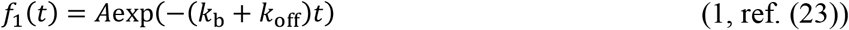

Observation times of genetically expressible fluorescent proteins are severely limited to the duration of a few acquisition frames due to photobleaching, limiting measurements of long-lived binding events (34). To extend observation times, the frame rate can be reduced by inserting a dark interval (τ_d_) after a short integration time (τ_int_). Scaling the photobleaching rate appropriately, Eq. 1 then becomes:

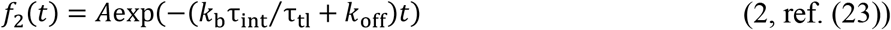

where the time-lapse time τ_tl_ is the sum of τ_int_ and τ_d_. The sum of two decay rates *k*_b_ and *k*_off_ can be approximated with an effective decay rate (*k*_eff_):

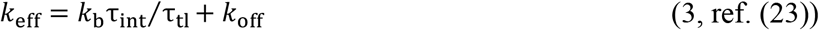

Rearrangement of Eq. 3 yields:

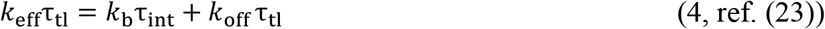

As *k*_b_τ_int_ is maintained constant at a certain imaging condition, *k*_eff_τ_tl_ increases linearly with τ_tl_, with the coefficient (slope) *k*_off_.

In systems with two sub-populations each dissociating at different rates *k*_off1_ and *k*_off2_, Eq. 2 then becomes:

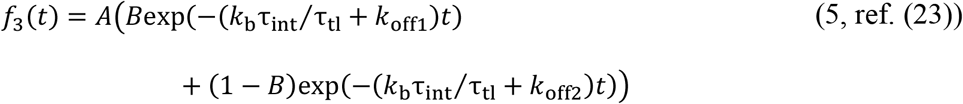

where *B* (0 < *B* < 1) and (1 – *B*) are the amplitudes of *k*_off1_ and *k*_off2_ sub-populations respectively. Similarly, a system with three kinetic sub-populations can be described by:

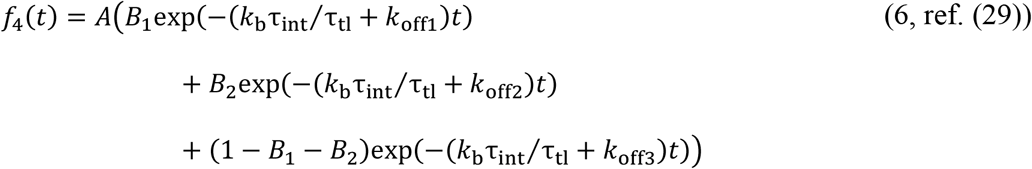

where *B*_1_, *B*_2_ (0 < *B*_1_, *B*_2_ < 1 and *B*_1_ + *B*_2_ < 1) and (1 – *B*_1_ – *B*_2_) represent the amplitudes of *k*_off1_, *k*_off2_ and *k*_off3_ sub-populations respectively.

### Experimental considerations

The specifics of the experimental setup for different model organisms should be tailored to requirements for the respective system. However, to provide the reader with a starting point, we describe the experimental configuration used in our lab to measure binding lifetimes of DNA-repair proteins labelled with the fluorescent protein, YPet in the model organism *Escherichia coli (E. coli)* (see Fig. S1 and ref. (32)). Bacterial cells in early exponential phase are loaded into a custom-built flow cell made up a glass coverslip and a quartz top. The bottom coverslip is functionalized with (3-Aminopropyl)triethoxysilane (APTES, Alfa Aesar, USA) to facilitate cell adhesion to the surface of the coverslip. The temperature of the flow cell is kept constant at 30 °C. Cells are supplied with aerated rich defined media (EZ rich defined medium supplemented with glucose, Teknova) to maintain fast growth. YPet is excited with 514-nm laser (Sapphire LP laser, Coherent, USA) in near-TIRF configuration (35) at a power density of 71 W/cm^2^ (measured directly above the inverted objective). Fluorescent signal is recorded using an electron-multiplying (EM)-CCD camera (Photometrics Evolve, Photometrics, USA), with an EM gain of 1,000. The camera exposure time is 0.1 s and time-lapse imaging is acquired with a 10-s τ_tl_ set (Table S2). Typically, a time-lapse imaging experiment lasts three to five hours and in generally four to ten experiments are required to obtain more than a thousand binding events at each τ_tl_.

Resolution of binding events in bacterial cells expressing copy numbers of fluorescent proteins in excess of ~ 20 copies per cell is challenging due to the limitations of particle tracking algorithms to resolve closely spaced foci. Further, distinguishing bound molecules from freely diffusive molecules in the cytosol is also challenging when copy numbers are high. In this case, to enable reliable observation of single-molecule cells are exposed to continuous illumination such that the majority of the emitters are darkened or photo-bleached, and only stochastically reactivated emitters are observed in single-molecule imaging conditions (36).

This setup allows us to unambiguously detect single-molecule foci using a relative signal-to-background ratio between six and eight. Foci detected in at least two consecutive frames within a 300-nm (3 pixels) radius are defined as a binding event. For each τ_tl_, all binding events are combined, and bootstrapping analysis is performed by randomly selecting with replacements 80% of all binding events. CRTDs are constructed from bootstrapped samples and are fit to exponential models to obtain *k*_eff_τ_tl_ plots, as well as *k*_b_ and τ.

### Simulating concurrent dissociation and photobleaching

In order to maintain full control of the kinetic variables, we chose to perform simulations of the experiment. Simulations of exponential distributions and curve-fitting were performed with custom-written program in MATLAB (The MathWorks, Natick, MA). We simulated exponential distributions (Eq. 2,5,6) using the *exprnd* function in MATLAB (Supplementary Notes). This function generates exponentially distributed random numbers with a specified decay constant. Here, each number returned by *exprnd* function represents the lifetime of a simulated ‘trajectory’. For the purposes of this work, we have not accounted for blinking of bound molecules that may yield prematurely truncated binding events. Accommodation of such a feature will require reasonable estimates of FP blinking under the conditions of the experiment that will be unique to the fluorescent probe used. To simulate a sub-population of molecules dissociating with a specified off rate, a set of trajectories was generated and binned to produce histograms with ten bins, whose edges correspond to frame times (integer multiples of τ_tl_). The *exprnd* function was iterated until the counts of the first bin exceeded the number of binding events in that sub-population (typically between three and six iterations, see Fig. S2). To simulate experiments where multiple sub-populations are present, each sub-population was simulated in defined proportions and all trajectories were pooled together. Finally, to generate the CRTDs, we rejected molecules in the first bin (0 to τ_tl_) and only carried forward observations from τ_tl_ to 10τ_tl_ to the next step in accordance with our definition of a binding event (or trajectories), *i.e,* the observation must be present in two consecutive frames.

To simulate uncertainty in each simulation sample, ten rounds of bootstrapping were performed, each involved randomly sampling 80% of the simulated population. Next, fitting was performed on each bootstrapped CRTDs (henceforth referred simply as CRTDs). First, the CRTD at each τ_tl_ was fit to a mono-exponential model to obtain *k*_eff_ (Eq. 2, 3 and Fig. 1C). These values for *k*_eff_, corresponding to the number of τ_tl_, were then used to construct the *k*_eff_τ_tl_ plot. Error bands in these plots represent standard deviations from ten bootstrapped samples.

Second, the CRTDs for all τ_tl_ were fit to objective functions based on Eq. 2, 5 and 6 (global fitting, see Supplementary Notes). The list of parameters, initial conditions, bound constraints, termination criteria and algorithm is presented in Table S1. Throughout the paper, *A* was set as a local parameter to mimic experimental conditions where counts may be different across τ_tl_, even though this often leads to less accurate results compared to when *A* was set as a global parameter (Fig. S3B-C).

For each simulation, outcomes from globally fitting the ten bootstrapped CRTDs were averaged and reported. To determine uncertainty in the estimate, we repeated the simulation a hundred times. The standard deviations of the binding lifetime (*σ*_τ_) from a hundred simulations using the same conditions was calculated according to Eq. 7.

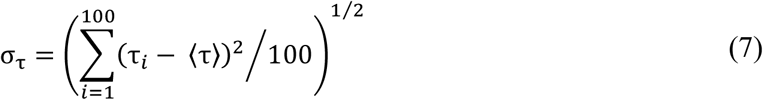

where <τ> denotes the true binding lifetime, which is calculated by 1/< *k*_off_>.

Unless otherwise stated, *k*_b_τ_int_ was fixed at 0.7 to mimic experimental values obtained in our published work (32). Four sets of τ_tl_ were used: 10-s τ_tl_, 100-s τ_tl_, the three- and five-τ_tl_ sets (Table S2).

## RESULTS

### Influence of experimental sample size on uncertainty of the estimate of the binding lifetime

First, we set out to investigate whether the size of the experimental data set influences the uncertainty in the error estimate of the outcomes from global fitting, such as the binding lifetime τ and photobleaching rate *k*_b_. This can be achieved by randomly selecting a fraction of experimental data (3%-30%) at each τ_tl_, following by bootstrapping and global fitting. Toward this goal, we revisited published data from our laboratory where interval imaging was used to determine dissociation kinetics of the transcription-repair coupling factor Mfd from DNA in live *E. coli* (32). The entire dataset (100%) contains between 1,000 to 2,000 trajectories (counts lasting at least two frames) at each τ_tl_ (Fig. 2A, right-most panel). Representative CRTDs following sub-sampling the experimental dataset (3%, 10% and 30%) at each τ_tl_ are shown in (Fig. 2A). While the *k*_eff_τ_tl_ plot derived from the whole dataset resembles a straight line, deviations from linear fits in *k*_eff_τ_tl_ plots can be seen when only a sub-set of experimental data was used (Fig. 2B).

**Figure 2.**
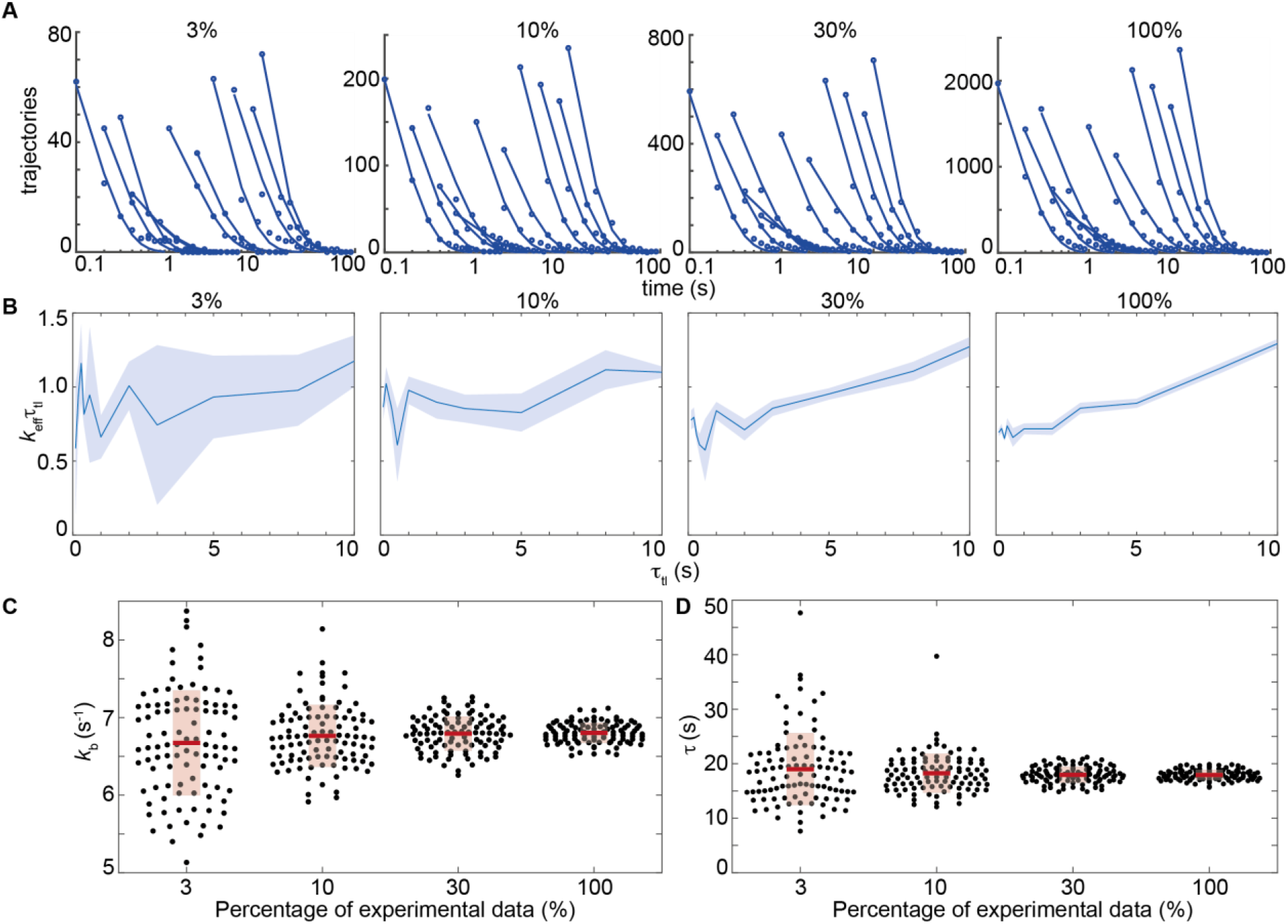
Determination of the photobleaching rate and binding lifetime from sub-sampling experimental data presented in ref (32). (A) Representative CRTDs when only 3%, 10% or 30% of experimental trajectories were randomly selected. Counts can be approximated as y-intercepts of exponential fits of CRTDs. The CRTD from the full dataset (right most panel) is reproduced from ref. (32). (B) *k*_eff_τ_tl_ plots of the corresponding CRTDs (above). Shaded error bands are standard deviations from ten bootstrapped samples. (C) Scatter plots show distributions of *k*_b_ obtained using global fitting 100 subsets of the experimental data at the indicated fraction. Each point represents the average of results from ten bootstrapped samples. (D) Scatter plots show distributions of τ obtained using global fitting 100 subsets of the experimental data at the indicated fraction. Similarly, each point represents the average of results from ten bootstrapped samples. Red bars and boxes represent means and standard deviations of the fitting outcomes of 100 subsets of the dataset respectively. The experimentally measured value of τ = 17.9 ± 0.9 s for the entire data set is reproduced from ref (32).

To determine the uncertainties in *k*_b_ and τ as a result of under-sampling, we repeated the subsampling a hundred times and *k*_b_ and τ values were obtained from global fitting using Eq. 2 (Fig. 2C-D). Here, uncertainties in the estimates of *k*_b_ and τ are smallest when the entire data set is used (2% and 5% respectively, Fig. 2C-D), and as expected, increase with decreasing number of counts (Fig. 2C-D). For *k*_b_, uncertainties increase from 3% to 10% as the percentage of experimental data drop from 30% to 3% while uncertainties in determining binding lifetimes increase from 8% to 35% (Fig. 2C-D).

Fitting individual CRTD to mono-exponential model to obtain *k*_eff_τ_tl_ plots has been suggested to be used as a guide to determine kinetic heterogeneity (23). Our analysis demonstrates that deviation from linear fits in the *k*_eff_τ_tl_ plots can potentially simply reflect under-sampling. Since deviations from linear fits in *k*_eff_Δ_tl_ plots can also be used to guide the choice of bi- and triexponential models (23), a fundamental question that faces users is, what governs the choice of exponential model? What is the minimum size of data, for which a multi-exponential model is appropriate for consideration? Are deviations in the *k*_eff_τ_tl_ plots reliable indicators for the choice of model? To explore these questions in greater detail, we chose to perform simulations that permit us to retain full control of the model parameters, and overcome practical limitations of generating large data sets from microscopy experiments.

### Case I: Influence of the size of the data set on the measured lifetime for a single dissociating species

We first explored the relationship between the number of counts (*n*) at each τ_tl_ and uncertainties in estimates of binding lifetimes from mono-exponential distributions. To this end, we simulated a population of molecules dissociating with *k*_off_ of 0.1 s^-1^, corresponding to a binding lifetime <τ> of 10 s, and photobleaching rate *k*_b_ of 7 s^-1^ (see Methods). While τ_int_ was constant at 0.1 s, τ_tl_ was varied from 0.1 s to 10 s (Table S2). These values of *k*_b_, <τ>, τ_int_ and τ_tl_ were initially chosen to closely match experimental values used in our published work (see Fig. 2 and ref. (32)). The theoretical *k*_eff_τ_tl_ plot is shown as the dashed line (Fig. 3A). At *n* = 1×10^3^ observations (Fig. 3A), the *k*_eff_τ_tl_ plot deviates noticeably from the theoretical line (purple curve). However, as *n* increases, the error bands reduce, and the plots closely resemble straight lines (purple curves, Fig. 3B-D). At 1×10^5^ observations, linearly fitting the *k*_eff_τ_tl_ plot (Fig. 3D) yielded a slope of 0.1 and y-intercept of 0.6992, reflecting the specified *k*_off_ (0.1 s^-1^) and *k*_b_τ_int_ (0.7). As expected, mono-exponential distributions with the same *k*_b_τ_int_ but smaller off rate (*k*_off_ = 0.01 s^-1^) or without off rate (*k*_off_ = 0 s^-1^) yielded lines with smaller slope (Fig. 3A-D, green curves) or essentially flat lines (Fig. 3A-D, black curves).

**FIGURE 3.**
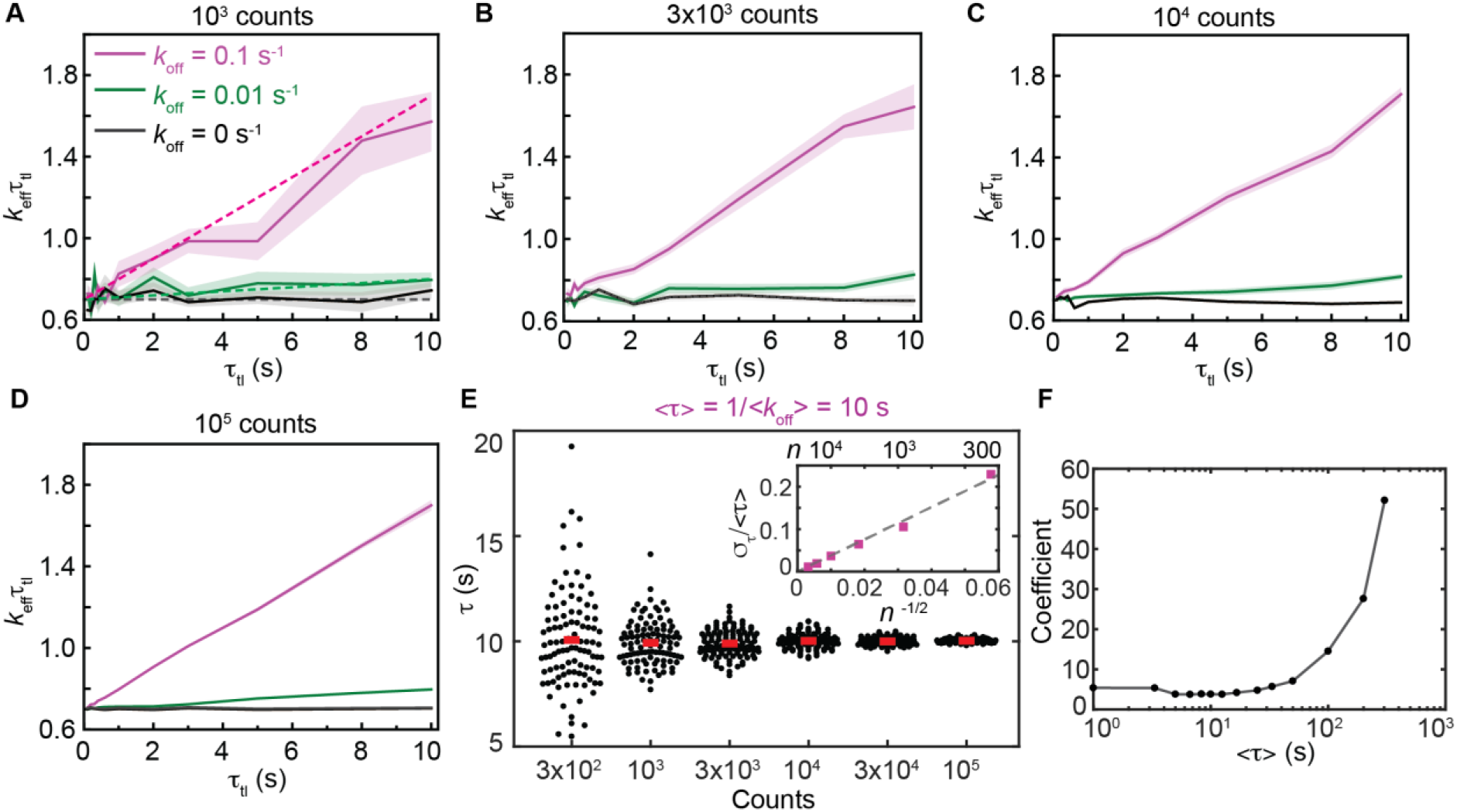
Determination of binding lifetimes from mono-exponential distributions. (A-D) *k*_eff_τ_tl_ plots of mono-exponential distributions with *k*bτint of 0.7 and *k*_off_ of 0.1 s^-1^ (purple curves), 0.01 s^-1^ (green curves) or 0 s^-1^ (black curves). Panels A-D reflect *k*_eff_τ_tl_ plots obtained from simulations containing number of observations (n) equaling (A) 1×10^3^, (B) 3×10^3^, (C) 1×10^4^ or (D) 1×10^5^ counts in the first bin (see Methods). (A) Dashed lines correspond to theoretical *k*_eff_τ_tl_ plots at the specified *k*_off_ values. Shaded error bands are standard deviations from ten bootstrapped samples. (E) Scatter plots show distributions of τ obtained using global fitting from 100 simulated samples for each *n* value. Red bars represent the mean values. (Inset) The relative error in determining τ (σ_τ_ /<τ>) reduces with *n*^-1/2^ for increasing *n*. Dashed line is the linear fit to six data points. (F) Coefficient in function of *σ*_τ_/<τ> versus *n*^-1/2^ at various <τ>. The sharp increase in coefficients for <τ> larger than 50 s indicates larger uncertainties in measuring slow processes when the maximum τ_tl_ is limited to 10 s.

To characterize the uncertainty (standard deviation, *σ*_τ_) in the estimate of the binding lifetime, we repeated the simulation a hundred times for each value of *n* and determined τ using global fitting (Fig. 3E). As expected for shot noise (37), the relative error *σ*_τ_/<τ> is proportional to the inverse of the square root of *n* with a coefficient of 3.8 (Fig. 3E, inset). Importantly, the coefficient fluctuates between 3.7 and 5.7 for <τ> ≤ 50 s, but rises sharply for <τ> greater than 50 s (Fig. 3F). This result demonstrates that the uncertainty in estimating lifetime of long-lived binding events becomes arbitrarily large when the extended lifetime of the fluorophore (by introduction of τ_d_) becomes comparable to the binding lifetime. In principle, this limit can be readily overcome by simply selecting larger τ_tl_ values; indeed, simulations of monoexponential distributions of long-lived binding events (τ = 100 s) indicated that *σ*_τ_ is lower at lower values of *n*, when τ_tl_ is extended to 100 s, compared to 10 s (Fig. S3).

Therefore, we propose that accurate measurements of lifetime of long-lived binding events require significant increases in either the number of observations (*n*) or the length of τ_tl_ for a fixed photobleaching rate. However, it should be noted that extension of τ_tl_ up to 100 s may not be experimentally feasible for all systems. In our work involving bacterial live-cell imaging in rich media, cell growth and division on the timescale of imaging limit the tracking binding events lasting on the timescale of tens of minutes. Practical limitations imposed by the model organism, growth conditions and choice of fluorescent protein dictate optimal experimental design.

Further, we anticipated that photobleaching rate also contributes to σ_τ_ as faster photobleaching reduces observation times. To examine the effect of *k*_b_τ_int_, we performed a comprehensive set of simulations with the 10-s τ_tl_ set (Table S2) and *k*_b_τ_int_ varying from 0.007 to 2.1 (*k*_b_ from 0.07 to 70 s^-1^ and τ_int_ from 0.01 to 0.1 s). We obtained the relationship between *σ*_τ_ /τ, *n* and *k*_b_τ_int_ as in Eq. 8.

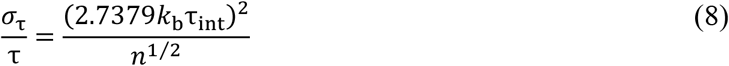

This formula describes the lower bound of errors as other sources of practical errors, such as localization uncertainties and experimental variations, have not been considered. The minimum number of observations required to determine τ (<τ> ≤ 50 s) with a given uncertainty is therefore:

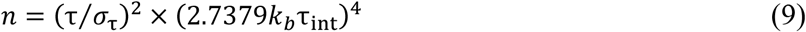

For example, when *k*_b_τ_int_ is 0.7, the number of observations required to achieve relative error of 10% in the estimate of τ (where <τ> ≤ 50 s) is about 1350 (see Fig. 3E). This equation also highlights the importance of using fluorophores with high photo-stability: a two-fold increase in *k*_b_ needs to be compensated by a 16-fold increase in *n*.

### Case II: Detection of two species with resolvable lifetimes

Next, we examined the situation where a second kinetic sub-population is present in the system. A second population with a faster off rate yields *k*_eff_τ_tl_ plots that deviate from straight lines (23). However, as we demonstrate deviations can also be a result of shot noise at low *n* (see Fig. 2B and Fig. 3A-D). To identify the minimum *n* at which one can determine with a specified confidence that a bi-exponential model is appropriate, we simulated CRTDs using Eq. 5. First, we performed simulations with off rates that are an order of magnitude apart: *k*_off1_ = 0.1 s^-1^ (intermediate rate) and *k*_off2_ = 1 s^-1^ (fast rate). The amplitude *B* of the intermediate dissociating population was varied from 10% to 90% (Fig. 4).

**FIGURE 4.**
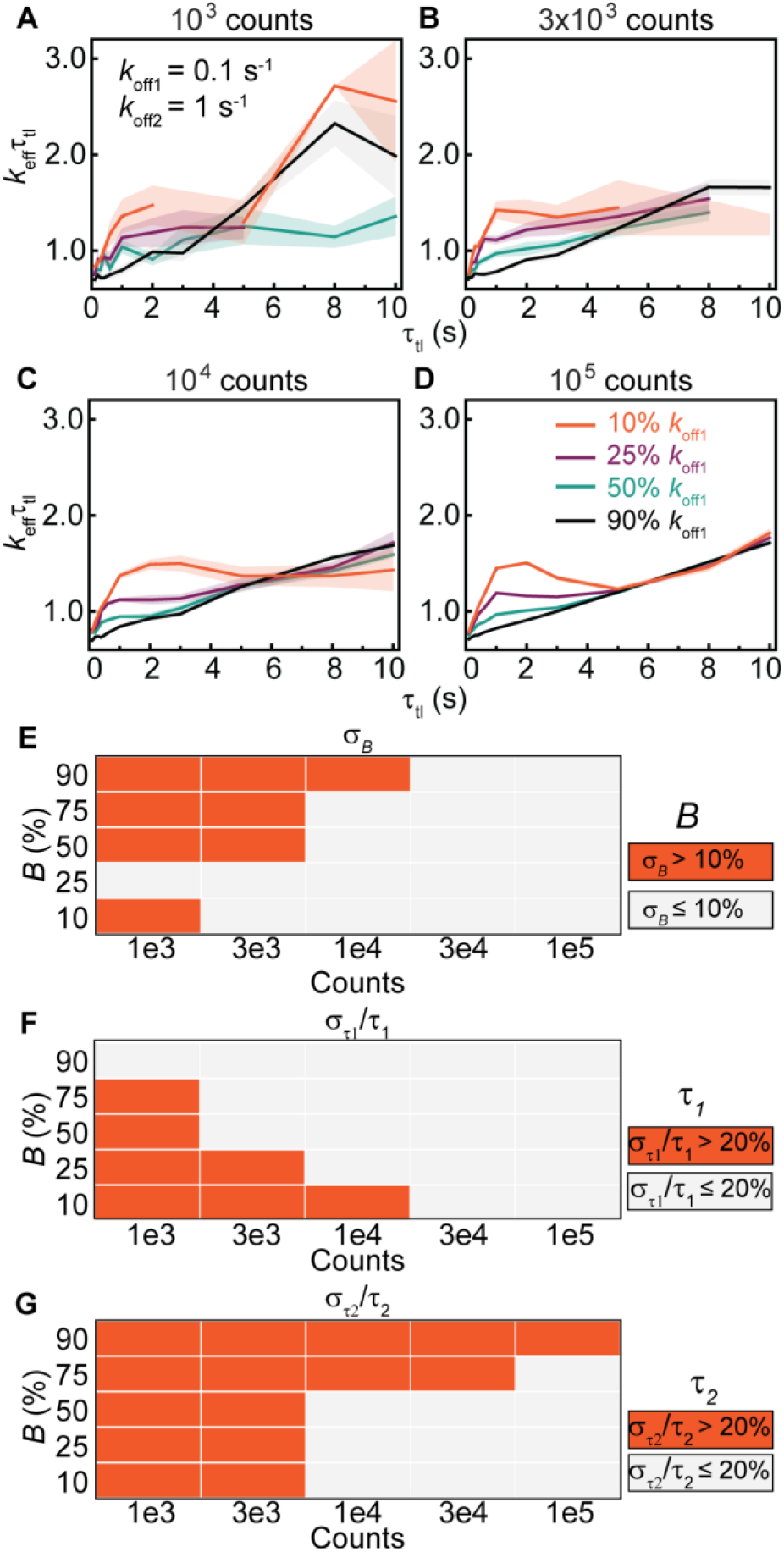
Determination of binding lifetimes and amplitudes from bi-exponential distributions with an intermediate rate (*k*_off1_) and a fast rate (*k*_off2_ = 10 *k*_off1_). (A-D) *k*_eff_τ_tl_ plots of bi-exponential distributions with *k*_b_τ_int_ of 0.7, *koff1* and *k*_off2_ of 0.1 and 1.0 s^-1^ respectively, with (A) 10^3^, (B) 3×10^3^, (C) 10^4^ or (D) 10^5^ observations. The amplitude of *k*_off1_ (B) is 10% (orange), 25% (purple), 50% (green) or 90% (black). Shaded error bars are standard deviations from ten bootstrapped samples. (E-G) Heatmaps show errors in estimates of B, τ_1_ and τ_2_ obtained using global fitting of 100 simulated distributions for each *n* value (see Fig. S4 for distributions).

When the majority of the population dissociates with the intermediate rate *k*_off1_ (B = 90%), the *k*_eff_τ_tl_ plots resemble those of mono-exponential distribution with the single *k*_off_ of 0.1 s^-1^ (compare Fig. 4A-D, black curves and Fig. 3A-D). As before, increasing the number of observations significantly improved the quality of the *k*_eff_τ_tl_ plots (Fig. 4A-D). These simulations reveal that a short-lived second sub-population does not manifest as a visible feature in the *k*_eff_τ_tl_ plots when it is present only to the extent of 10% in the observations. To examine if the two populations could be resolved with global fitting using the bi-exponential model, we determined binding lifetimes and amplitudes from 100 simulations (Fig. S4). Unsurprisingly, we found that the accuracies and precisions of determining B, τ_1_ and τ_2_ increase with n. While estimation of τ_1_ is robust (Fig. 4F, Fig. S4B), global fitting of CRTDs to the biexponential model at low counts suffers from a bias towards the fast dissociating subpopulation, with its amplitude being overestimated and τ_2_ being underestimated (Fig. 4E, G, Fig. S4A, C). This bias is observed to a lesser extent when *k*_off1_ is present at 75% or 50% (Fig. 4E-G, Fig. S4).

As the amplitude of the fast dissociating sub-population increased (B equal to 25% or 10%), fewer observations were found at long intervals. Insufficient counts resulted in missing data points at these τ_tl_ (τ_tl_ ≥ 5 s) in *k*_eff_τ_tl_ plots at low counts (10^3^ and 3×10^3^, Fig. 4A-B). However, the *k*_eff_τ_tl_ plots extended to the full τ_tl_ range of 10 s when *n* increases to 10^4^ and 10^5^ (Fig. 4C-D). As expected, deviations from straight lines were found in the 0-5 s regime, reflecting the presence of the fast dissociating sub-population. Since contributions from the fast dissociating sub-population drop sharply at long timescales, the *k*_eff_τ_tl_ plots converge to the straight line exhibited by mono-exponential distributions with *k*_off1_ (Fig. 4C-D). Further analysis by integrating the area under the peaks in the 0-to-5 s region shows the area increases exponentially with the amplitude of the fast dissociating sub-population (Fig. S5). When the fast dissociating sub-population represents the majority, the accuracy and precision in determining B, τ_1_ and τ_2_ also increase with *n* (Fig. 4E-G, Fig. S4).

Based on the observation that accurate measurements of long-lived binding events require the extension of τ_tl_ to greater than 10 s, we anticipated that resolving two kinetic sub-populations [one with a slow rate (*k*_off1_ of 0.01 s^-1^; <τ_i_> = 100 s) and an intermediate rate (*k*_off2_ of 0.1 s^-1^; <τ_2_> = 10 s)] is challenging when the largest τ_tl_ is 10-s. Consistent with this, the *k*_eff_τ_tl_ plots in the 0-10s range appear linear (Fig. 5A-C), resembling those of mono-exponential distributions. Hence, we attempted to fit the CRTDs at 10^5^ counts to mono-exponential model (Eq. 2), yielding apparent binding lifetimes (τ*) that lie between <τ_1_> and <τ_2_> (Fig. 5D). Fitting mean τ* vs. *B* to exponential function results in Equation 10:

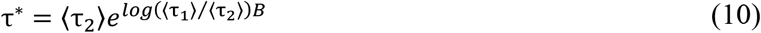

**FIGURE 5.**
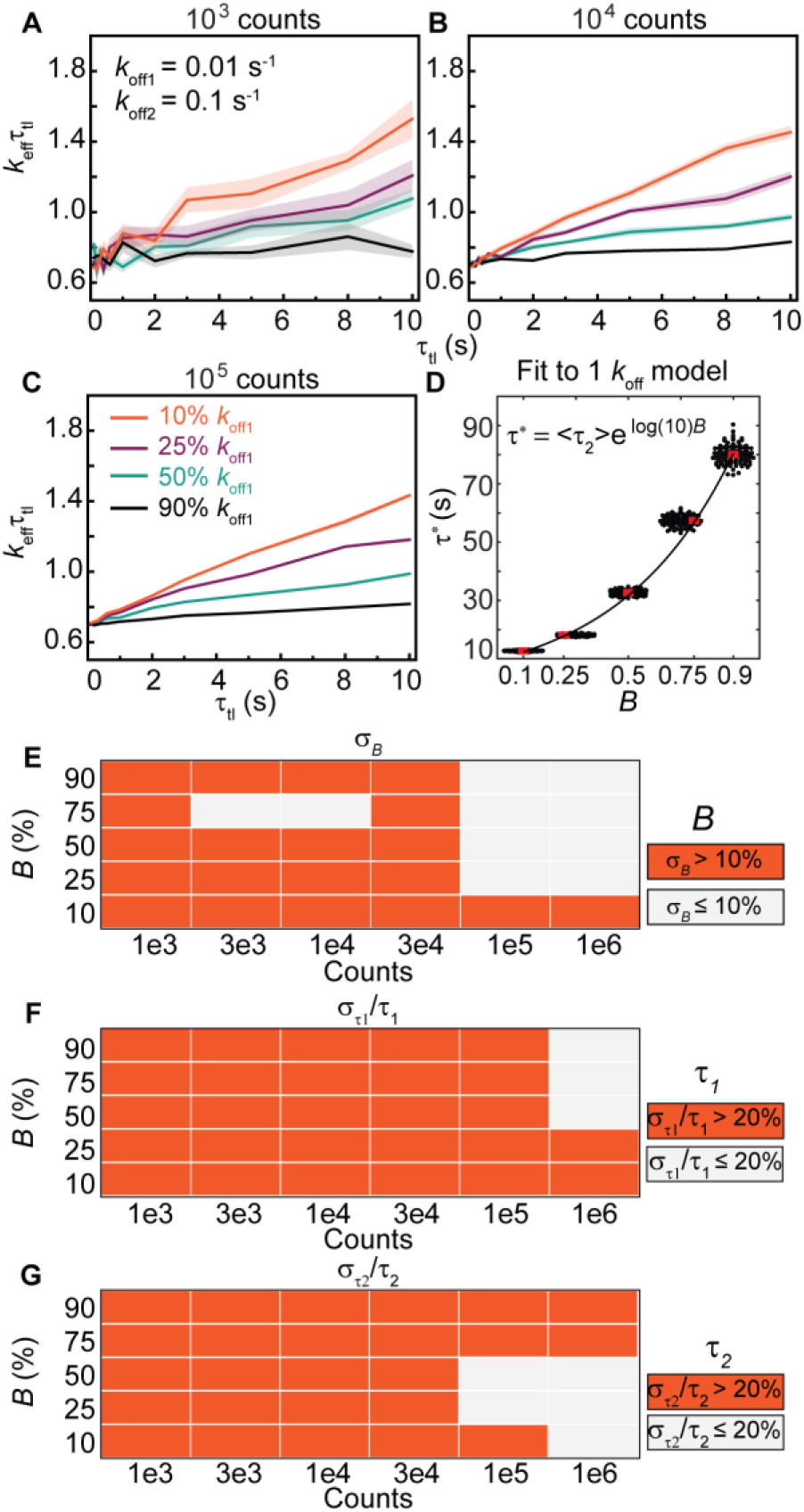
Determination of binding lifetimes and amplitudes from bi-exponential distributions with a slow rate (*k*_off1_) and an intermediate rate (*k*_off2_ = 10*k*_off1_). (A-C) *k*_eff_τ_tl_ plots of bi-exponential distributions with *k*_b_τ_int_ of 0.7, *k*_off1_ and *k*_off2_ of 0.01 and 0.1 s^-1^ respectively, with (A) 10^3^, (B) 10^4^ or (C) 10^5^ observations. The amplitude of *k*_off1_ (B) is 10% (orange), 25% (purple), 50% (green) or 90% (black). Shaded error bars are standard deviations from ten bootstrapped samples. (D) Scatter plots show distribution of apparent τ (τ*) obtained from fitting of 100 simulated bi-exponential distributions at a specified *B* and 10^5^ counts to monoexponential model. Line is exponential fit between the average of τ* (red bars) and B. (E-G) Heatmaps show errors in estimates of B, τ_1_ and τ_2_ obtained using global fitting of 100 simulated distributions for each *n* value (see Fig. S6 for distributions).

Thus, *B* can be derived from τ* where <τ_1_> and <τ_2_> are known.

From the simulations, fitting the CRTDs with *n* less than 3×10^4^ to the bi-exponential model yields unreliable results (Fig. 5E-G, Fig. S6). Across various amplitudes of *k*_off1_, the species with lifetime τ_1_, is often underestimated and corresponds to τ* at that amplitude (compare Fig. 5F to Fig. 5D). Similarly, τ_2_ is also underestimated, but eventually approaches <τ_2_> of 10 s when *n* reached 10^6^ counts and the amplitude of *k*_off2_ sub-population is more than 25% (Fig. 5G).

On the other hand, when the above distributions were simulated using the 100-s τ_tl_ set, deviations from straight lines in *k*_eff_τ_tl_ plots were observed in the 0-30s regime and when *B* is smaller than 75% (Fig. S7A). In this case, as expected, accuracies in determining B, τ_1_ and τ_2_ follow the same trends as discussed in Fig. 4 (Fig. S7B-D).

### Case III: Detection of two species with closely matched lifetimes

Due to the resolution limit that is inherent to exponential analysis (34), we anticipated the ability to resolve rates that are closely spaced would reduce. To test this hypothesis, we simulated bi-exponential distributions with rates that are only three-fold apart: an intermediate rate *k*_off1_ of 0.1 s^-1^ and a fast rate *k*_off2_ of 0.3 s^-1^. Under conditions that yield sufficient observations at long intervals (*n* ≥ 10^4^), examination of the *k*_eff_τ_tl_ plots often fails to identify the presence of multiple sub-populations in the form of deviation from straight lines (Fig. 6A-D). Only when the fast rate is present at 90%, can deviations be observed in the form of a broad convex spanning from 0 to 10 s (orange curves, Fig. 6C-D). Fitting to Eq. 5 yields unreliable results for *B* and τ_2_ for *n* ≤ 10^4^ (Fig. 6E, G, Fig. S8A, C) whereas the accuracy in determining τ_1_ requires 3 x10^3^ observations or <B> to be larger than 25% (Fig. 6F, Fig. S8B). Fitting CRTDs at low counts (*n* ≤ 10^4^) to the bi-exponential model should be avoided as one often obtains two kinetics sub-populations with artificially enhanced rate separation and substantial amplitudes, regardless of the true amplitudes (Fig. 6E-G, Fig. S8).

**FIGURE 6.**
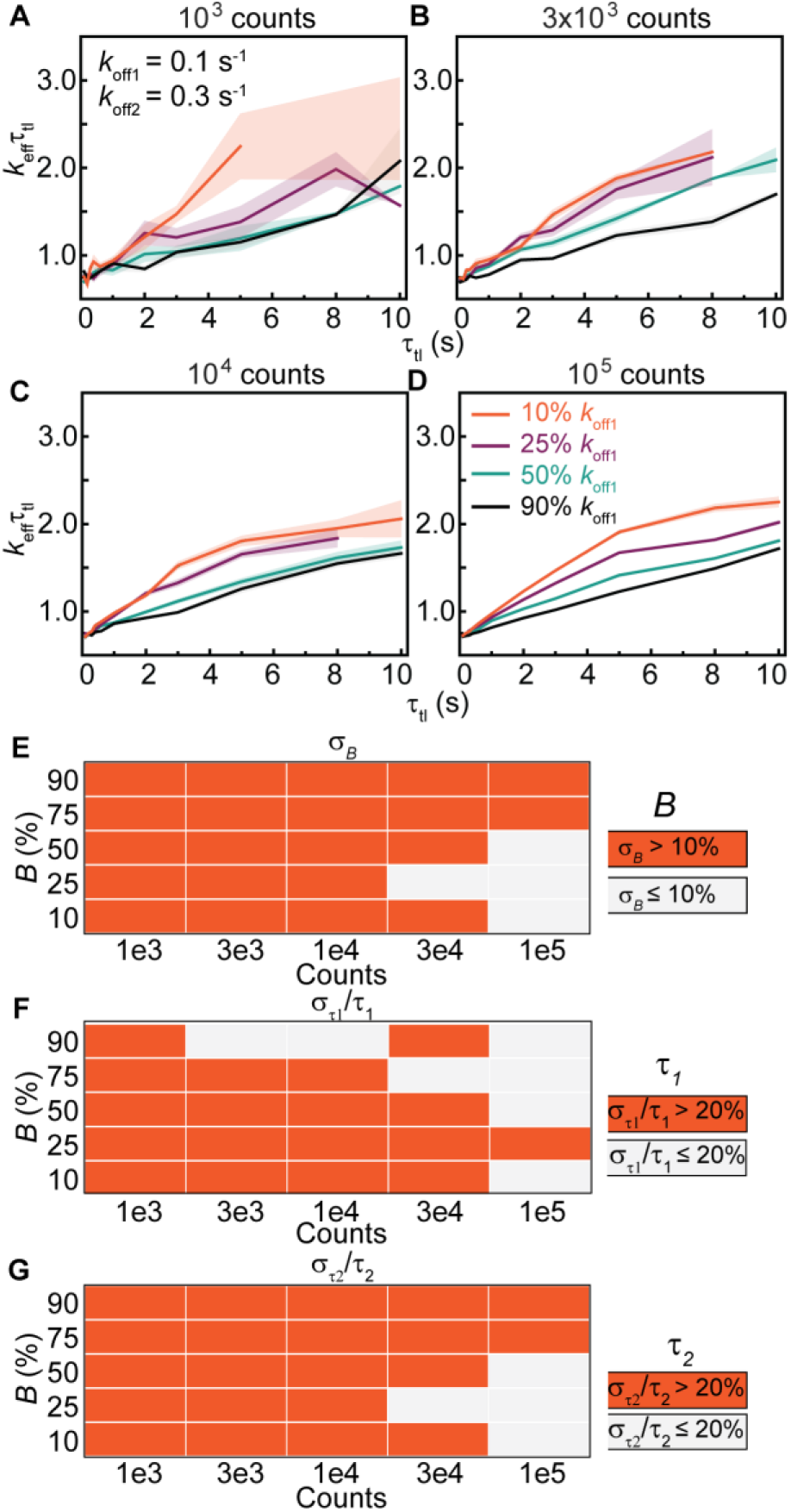
Determination of binding lifetimes and amplitudes from bi-exponential distributions with closely spaced rates (*k*_off2_ = 3 *k*_off1_). (A-D) *k*_eff_τ_tl_ plots of bi-exponential distributions with *k*_b_τ_int_ of 0.7, *k*_off1_ and *k*_off2_ of 0.1 and 0.3 s^-1^ respectively, with (A) 10^3^, (B) 3×10^3^, (C) 10^4^ or (D) 10^5^ observations. The amplitude of *k*_off1_ (B) is 10% (orange), 25% (purple), 50% (green) or 90% (black). Shaded error bars are standard deviations from ten bootstrapped samples. (E-G) Heatmaps show errors in estimates of B, τ_1_ and τ_2_ obtained using global fitting of 100 simulated distributions for each *n* value (see Fig. S8 for distributions).

### Case IV: Detection of three species

The resolution limit as well as dynamic range limit that we demonstrated above raise the question if tri-exponential distributions can be faithfully resolved under the specified experimental condition (ranges of τ_tl_ and n). To address this issue, we simulated tri-exponential distributions (Eq. 6), with off rates spanning two orders of magnitude (0.01, 0.1 and 1 s^-1^), using the 100-s τ_tl_ set. The diversity in *k*_eff_τ_tl_ plots obtained by varying *B*_1_ and *B*_2_ is illustrated in Fig. 7A. Three kinetic sub-populations are apparent when B1 is a third of *B*_2_ and *B*_2_ in turn is a third of *B*_3_ (1 – *B*_1_ – *B*_2_). We further characterized uncertainties in amplitudes and binding lifetimes obtained using global fitting to the tri-exponential model (Methods). In general, accuracy in determining the amplitudes and lifetimes improves with increasing *n* (Fig. 7B-F, Fig. S9). However, when the slowly dissociating sub-population dominates (*B*_1_ = 9/13), increasing *n* does not yield more accurate estimates. As in the case of the bi-exponential simulations, we observed consistent biases towards faster binding lifetimes (Fig. S9).

**FIGURE 7.**
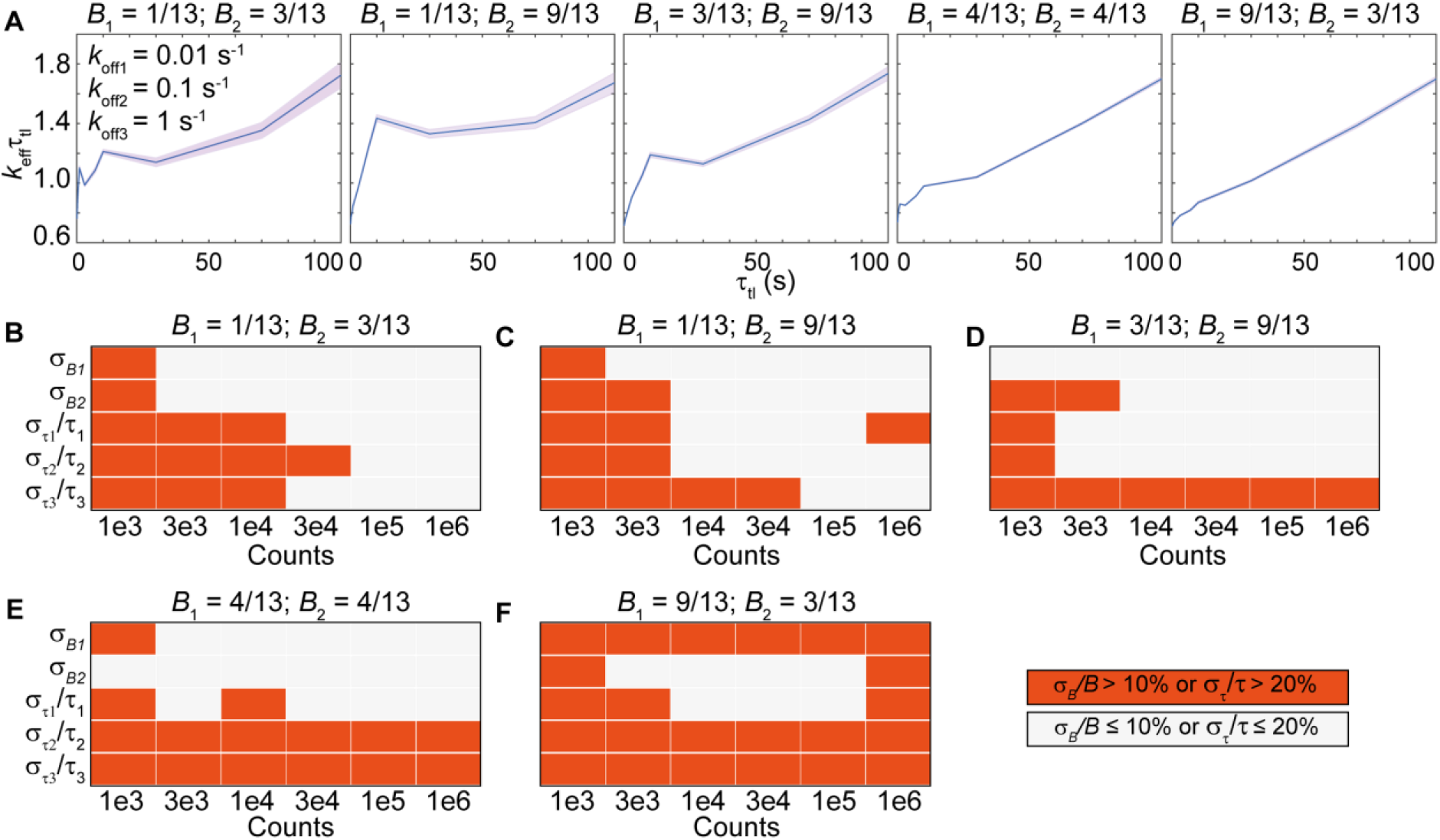
Determination of binding lifetimes and amplitudes from tri-exponential distributions with a slow rate (*k*_off1_), an intermediate rate (*k*_off2_ = 10*k*_off1_) and a fast rate (*k*_off3_ = 10*k*_off2_), using the 100-s τ_tl_ set. From left to right, five panels in each row correspond to different amplitudes of each sub-population (displayed on top). (A) *k*_eff_τ_tl_ plots of tri-exponential distributions with *k*_b_τ_int_ of 0.7, *k*_off1_, *k*_off2_ and *k*_off3_ of 0.01, 0.1 and 1 s^-1^ respectively, with 10^6^ observations. Shaded error bands are standard deviations from ten bootstrapped samples. (B-F) Heatmaps show errors in estimates of *B*_1_, *B*_2_, τ_1_, τ_2_ and τ_3_ obtained using global fitting of 100 simulated distributions at various pre-set values of *B*_1_ and *B*_2_ (see Fig. S9 for distributions).

### The choice of τ_tl_

Given a finite amount of experimental time, should one sample with more τ_tl_ values (increase interval) or obtain more observations (increase *n*) with a set containing fewer τ_tl_ values? To identify the optimum choice of τ_tl_, we simulated bi-exponential distributions with an intermediate rate (*k*_off1_ = 0.1 s^-1^) and a fast rate (*k*_off2_ = 1 s^-1^) using a τ_tl_ set containing either three (*N*_3_) or five (*N*_5_) τ_tl_ values, ranging from 0.1 to 10 s (Table S2). Since fitting outcomes are unreliable in the three τ_tl_ set (compare Fig. S10 to Fig. S11), we decided to examine the simulations with the five τ_tl_ set further. These simulations yielded *k*_eff_τ_tl_ plots that closely resemble those in Fig. 4 (see Fig. 8A-D) and similarly, deviations from straight lines are also reliable indicators of kinetic heterogeneity when *B* is less than 90%. As expected, estimates of B, τ_1_ and τ_2_ are more accurate with larger *n* (Fig. S10).

**FIGURE 8.**
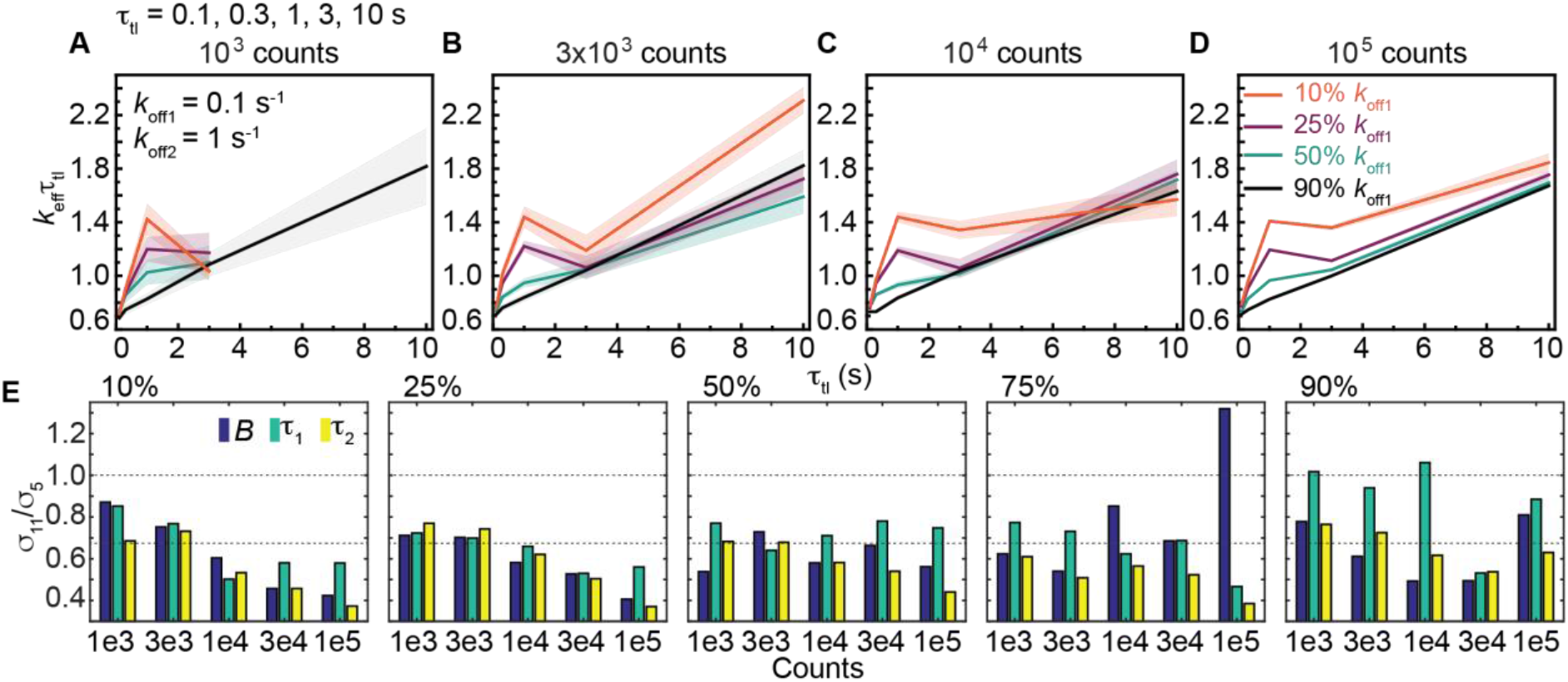
Determination of binding lifetimes and amplitudes from bi-exponential distributions using a τ_tl_ set containing five τ_tl_ values (Table S2), an intermediate rate (*k*_off1_ = 0.1 s^-1^) and a fast rate (*k*_off2_ = 1 s^-1^). (A-D) *k*_eff_τ_tl_ plots of bi-exponential distributions with (A) 10^3^, (B) 3×10^3^, (C) 10^4^ or (D) 10^5^ observations. The amplitude of *k*_off1_ (B) is 10% (orange), 25% (purple), 50% (green) or 90% (black). Shaded error bands are standard deviations from ten bootstrapped samples. (E) Bar plots show ratios of error estimates obtained from simulations with eleven and five τ_tl_ values at the same *n*. Blue: *B*, green: τ_1_, yellow: τ_2_.

Compared the simulations using the five τ_tl_ and the 10-s τ_tl_ (11 τ_tl_ values) sets for the same *n*, errors of estimates are almost always smaller in simulated distributions with the 10-s τ_tl_ set (*σ*_11_/*σ*_5_ < 1, see Fig. 8E). By extension of Eq. 8, error ratios (*σ*_11_/*σ*_5_) smaller than 1√V(11/5) or 0. 67 indicate the benefit of increasing Ninterval outweighs the benefit of increasing *n* with the five τ_tl_ set whereas error ratios larger than 0.67 represent redundancy in τ_tl_. Redundancy in τ_tl_ was observed in some cases when the intermediate dissociating sub-population is the majority (B between 75% and 90%) (Fig. 8E). However, when the majority dissociates with the fast rate (B between 10% and 50%), the benefit of sampling with more τ_tl_ is clear (σ_11_/σ_5_ < 0.67), especially with *n* ≥ 10^4^. Thus, we concluded the net benefit of increasing Ninterval is greater than increasing the number of counts with a set of fewer τ_tl_ values.

## DISCUSSION

In this work, we used experimental and simulated data to explore the influence of shot noise, resolution limit and dynamic range limit on resolving multiple kinetic sub-populations in single-molecule time-lapse imaging experiments (Fig. 9). Within the dynamic range and resolution limit, determination of binding lifetimes and amplitudes in mono-exponential and multi-exponential distributions are reliable in general, especially with at least 10^4^ counts.

**FIGURE 9.**
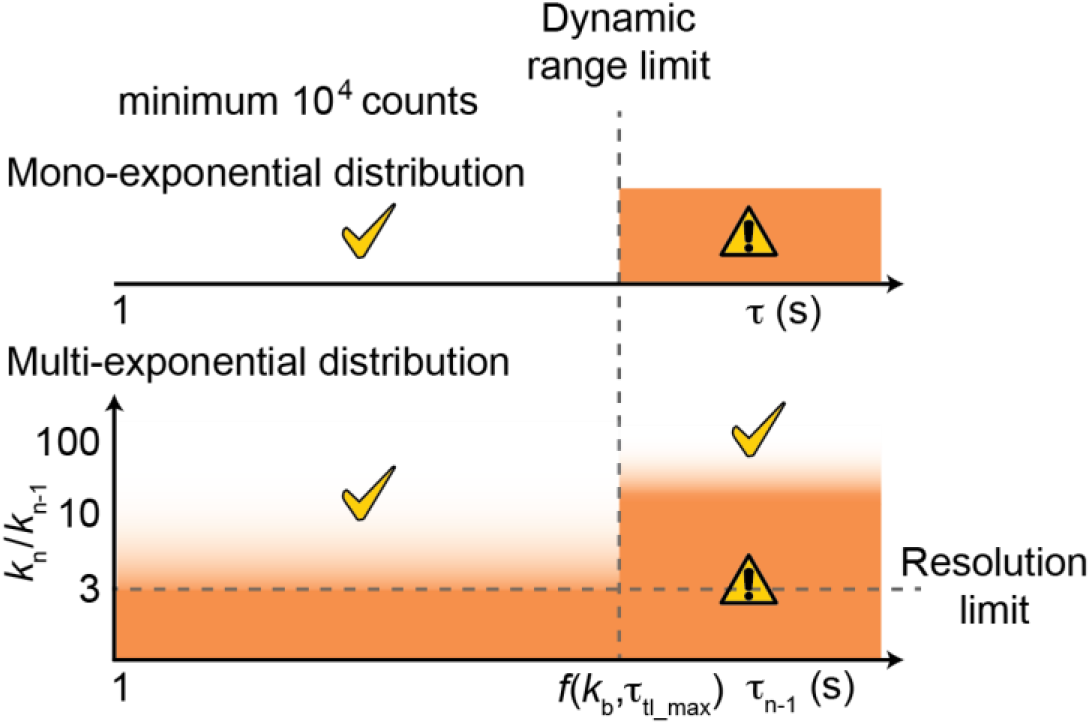
Dynamic range and resolution limits in resolving multiple populations using the time-lapse imaging technique with photobleaching-prone fluorescent probes. Dynamic range limit is a function of photobleaching rate (*k*_b_) and the maximum τ_tl_ (τ_tl_max_) used in experimental conditions. τ_n-1_ is the longest binding lifetime in a multi-exponential distribution within pairs of off rates *k*_n_ and *k*_n-1_. Orange zones indicate conditions where errors in estimates of τ and the amplitude are high.

As showed in Eq. 8, the relative error in τ determination scales with the square of *k*_b_τ_int_ and the inverse square root of *n*. This emphasizes the importance of choosing imaging conditions to minimize *k*_b_τ_int_ as a two-fold increase in *k*_b_τ_int_ needs to be compensated by a 16-fold increase in *n*. A balance has to be struck here to ensure good signal-to-background ratio, a prerequisite for reliable particle tracking. These findings also highlight the importance of developing and using fluorophores with higher photo-stability and brightness for live-cell applications as these would greatly reduce uncertainties in measurements. In practice, the choice of fluorescent protein should be made with great care, as fluorescent proteins often exhibit undesirable properties that limit their utility (38–42).

Errors obtained from repeating the experiments can be an underestimation compared to inherent errors conferred by shot noise when fitting is ill-conditioned (43), which is often the case when minimizing using multi-objective functions (44). Therefore, reports of binding lifetimes measurements using these time-lapse imaging approaches should clearly state *k*_b_τ_int_ from fitting and *n* from experimental data. This would enable a theoretical error estimation of τ and avoid over-interpretation of experimental results.

We found *k*_eff_τ_tl_ plots useful for guiding the fitting model when the number of counts is sufficiently large (more than 10^4^) as deviations from straight lines faithfully reflect heterogeneity in binding kinetics. The reverse is not necessarily true. Good linear fits, seen at large *n* values, can reflect one of the following three scenarios: (i) the absence of multiple populations, (ii) sub-populations with off rates that are within the resolution limit, or (iii) subpopulations where the off rate of one population lies beyond the dynamic range. This dynamic range is determined by the photobleaching rate and the maximum τ_tl_ used in the experiment. When the mono-exponential model is used to fit those data, an apparent binding lifetime τ*, whose value lies between the two true binding lifetimes, is obtained. While sub-optimal, τ* depends on the proportion of molecules in each kinetic sub-population: a larger presence of the fast dissociating sub-population yields smaller τ*. This in turn can report on change in binding kinetics when the biology is manipulable – for instance with binding partners or drugs.

Can statistical information such as reduced *χ*^2^ be used to decide the model that best describes the data? Computing these criteria requires the determination of the degree of freedom, which still needs to be analytically derived for the non-linear models used in this method (45–48). Instead of using statistical criteria, the selection of the fitting model using *k*_eff_τ_tl_ plot can be complemented with experimental design. For example, in case where a bi-exponential model is invoked, it might be tempting to attribute sub-populations to molecules performing certain activities such as binding of DNA repair proteins to a damaged or non-damaged substrate. These hypotheses can be tested using structure-function mutants in which one or few catalytic activities are inhibited, hence, yielding predictable changes in *k*_eff_τ_tl_ plots and fitting results. Finally, where possible, we recommend approaches that utilize multiple experimental designs to reproducibly observe or enrich the hypothesized populations.

## SOFTWARE

Our algorithms are freely available as open source MATLAB codes from https://github.com/hanngocho/off-rate-simulation.

## SUPPORTING INFORMATION

The supplementary information is available following publication.

## AUTHOR CONTRIBUTIONS

H. N.H., H.G. and A.M.v.O. designed research. H.N.H. wrote the codes, conducted the simulations and wrote the first draft. D.Z. contributed to the codes, with support from J.K. H.G. and A.M.v.O revised the manuscript and supervised research.

## ACKOWLEDGEMENTS

This work was supported by Australian Research Council Grant DP180100858, Australian Laureate Fellowship FL140100027 (to A.M.v.O.). We thank Dr. Joris M. H. Goudsmits for assisting with the simulation codes in MATLAB. D.Z. gratefully acknowledges support from the Elite Network of Bavaria (ENB) program “Macromolecular Science”.

