## Supplementary Information for "Identification of multiple kinetic populations of DNA-binding proteins in live cells"

**Running title: Measuring binding lifetimes in cells**

Han N. Ho[1,2,6], Daniel Zalami[3], Jürgen Köhler[3,4,5], Antoine M. van Oijen[1,2,*] and Harshad Ghodke[1,2,*]

[1]Molecular Horizons and School of Chemistry and Molecular Bioscience, University of Wollongong, Wollongong, Australia

[2]Illawarra Health and Medical Research Institute, Wollongong, Australia

[3]Spectroscopy of soft Matter, University of Bayreuth, Bayreuth, Germany

[4]Bavarian Polymer Institute, Bayreuth, Germany

[5]Bayreuth Institute of Macromolecular Research (BIMF), University of Bayreuth, Bayreuth, Germany

[6]Present address: The Francis Crick Institute, London, UK

### Supplementary table

**Table S1.** Initial conditions, constraints and termination tolerance used in global fitting. $n_0$ is the minimum number of counts in the second bin across $\tau_{tl}$.

| Model | Initial conditions | Bound constraints | Termination tolerance | Algorithm | MATLAB function |
|---|---|---|---|---|---|
| **Mono (Eq. 2)** | $k_b = 1$ s$^{-1}$<br>$k_{off} = 1$ s$^{-1}$ | $k_b > 0$ s$^{-1}$<br>$0$ s$^{-1} < k_{off} < 1/\tau_{int}$ s$^{-1}$ | $10^{-6}$ | trust-region-reflective | *lsqnonlin* |
| **Bi (Eq. 5)** | $k_b = 1$ s$^{-1}$<br>$k_{off1} = 1$ s$^{-1}$<br>$B = 0.5$<br>$k_{off2} = 2$ s$^{-1}$ | $k_b > 0$<br>$10^{-3}$ s$^{-1} < k_{off1} < 1/\tau_{int}$ s$^{-1}$<br>$1/n_0 < B < 1 - 1/n_0$<br>$10^{-3}$ s$^{-1} < k_{off2} < 1/\tau_{int}$ s$^{-1}$ | $10^{-6}$ | trust-region-reflective | *lsqnonlin* |
| **Tri (Eq. 6)** | $k_b = 1$ s$^{-1}$<br>$k_{off1} = 0.05$ s$^{-1}$<br>$B_1 = 0.3$<br>$k_{off2} = 0.5$ s$^{-1}$<br>$B_2 = 0.3$<br>$k_{off2} = 5$ s$^{-1}$ | $k_b > 0$ s$^{-1}$<br>$10^{-3}$ s$^{-1} < k_{off1} < 1/\tau_{int}$ s$^{-1}$<br>$1/n_0 < B_1 < 1 - 1/n_0$<br>$10^{-3}$ s$^{-1} < k_{off2} < 1/\tau_{int}$ s$^{-1}$<br>$1/n_0 < B_2 < 1 - 1/n_0$<br>$10^{-3}$ s$^{-1} < k_{off3} < 1/\tau_{int}$ s$^{-1}$<br>$B_1 + B_2 < 1 - 2/n_0$ | $10^{-9}$ | trust-region-reflective | *fmincon* |

**Table S2.** The $\tau_{tl}$ sets used in the study.

| $\tau_{tl}$ **sets** | $\tau_{tl}$ **values (s)** |
|---|---|
| 10-s | 0.1, 0.2, 0.3, 0.4, 0.6, 1, 2, 3, 5, 8, 10 |
| 100-s | 0.1, 0.3, 0.7, 1, 3, 7, 10, 30, 70, 100 |
| Three- | 0.1, 1, 10 |
| Five- | 0.1, 0.3, 1, 3, 10 |

### Supplementary figures

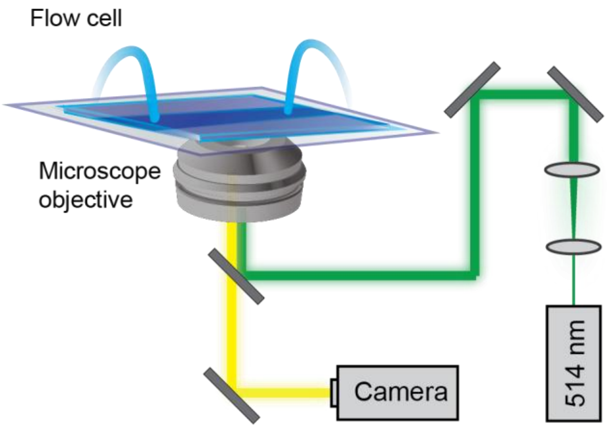


**Figure S1. Schematic of experimental setups in single-molecule live-cell imaging.** Bacteria expressing fluorescently labelled proteins are loaded in a flow cell with a constant supply of media at 30 ºC. The fluorescent label (YPet) is excited with 514-nm light and fluorescence signal is recorded with an electron-multiplying CCD camera.

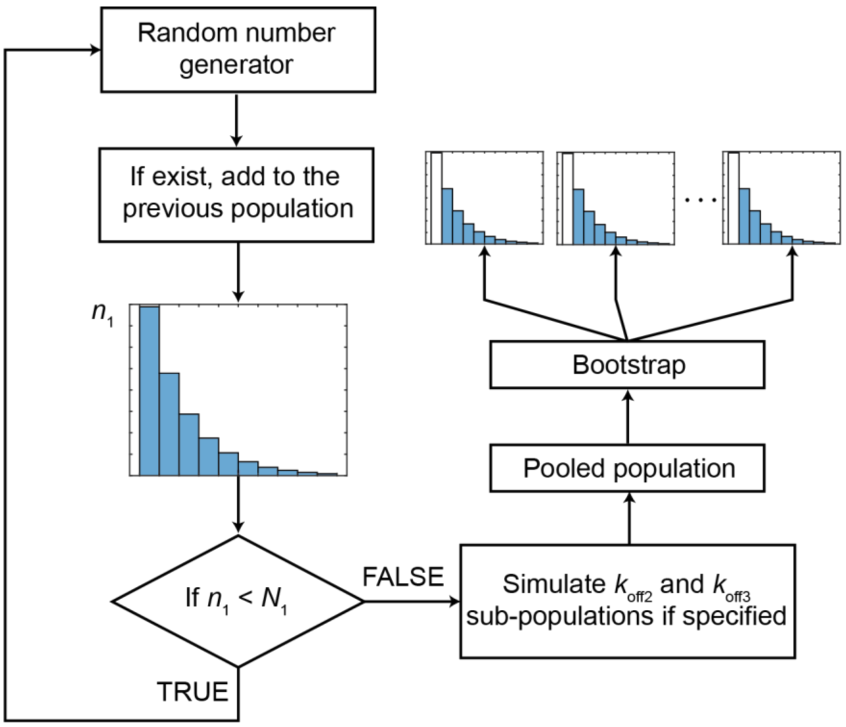


**Figure S2. Schematic of the simulation of the cumulative residence time distribution (CRTD) at a specified $\tau_{tl}$.** The molecules were generated by a random number generator to produce a group of numbers following an exponential distribution (defined by $k_{off1}$, $k_b$, $\tau_{int}$ and $\tau_{tl}$) (see Eq. 4-6 in main text). The number generator function was called a few times (typically 3-6) until the number of molecules in the first bin ($n_1$) of the histogram exceeded the user-specified number of molecules ($N_1$, $N_1 = A \times B$ in mono-exponential distribution, or $N_1 = A \times B_1$ in multiple-exponential distribution). The $k_{off2}$ and $k_{off3}$ sub-populations were simulated in the same manner. Then, molecules from all simulated sub-populations were pooled and subject to bootstrapping analysis to construct the bootstrapped CRTDs (referred simply as CRTDs). This procedure was repeated for all specified values of $\tau_{tl}$. The global fitting was performed on CRTDs from all $\tau_{tl}$, using a CRTD for each $\tau_{tl}$.

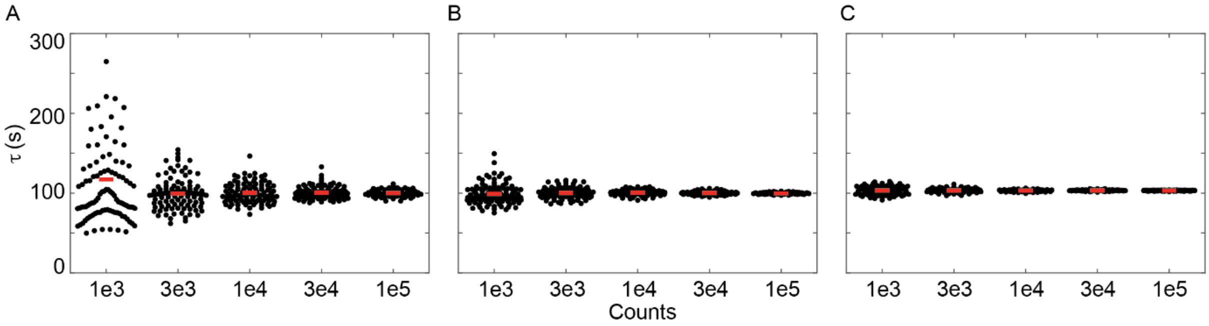


**Figure S3.** Scatter plots show distributions of $\tau$ obtained using global fitting on 100 simulated mono-exponential ($<\tau>$ = 100 s) for each *n* value. (A) Simulation using the 10-s $\tau_{tl}$ set. (B) Simulation using the 100-s $\tau_{tl}$ set. (C) Simulated data from (B) were globally fitted with the amplitude as the global parameter. Apart from this panel, all global fittings in this study were performed with *A* as the local parameter. Red bars represent the averages.

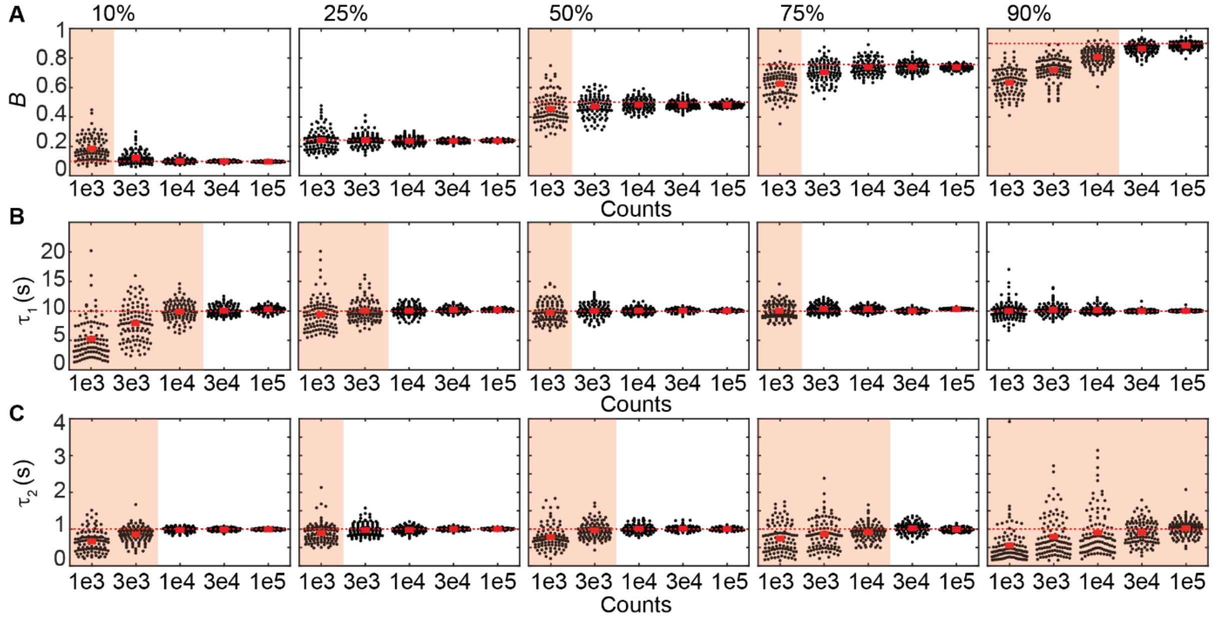

**Figure S4.** Determination of time constants and amplitudes from bi-exponential distributions with an intermediate rate ($k_{off1}$) and a fast rate ($k_{off2} = 10k_{off1}$). (A-C) Scatter plots show distributions of $B$, $\tau_1$ and $\tau_2$ obtained using global fitting from 100 simulated distributions for each $n$ value. Each panel corresponds to a pre-set $B$, which increases from 10%, 25%, 50%, 75% to 90% from left to right. In each panel, $n$ increases from $10^3$ (1e3) to $10^5$ (1e5). Dashed lines and red bars represent the true values and the average respectively. Orange shades represent distributions where $\sigma_B$ is larger than 0.1 or $\sigma_\tau/\tau$ is larger than 20%. To enhance visibility, outliers (less than 5% when present) were omitted from scatter plots.

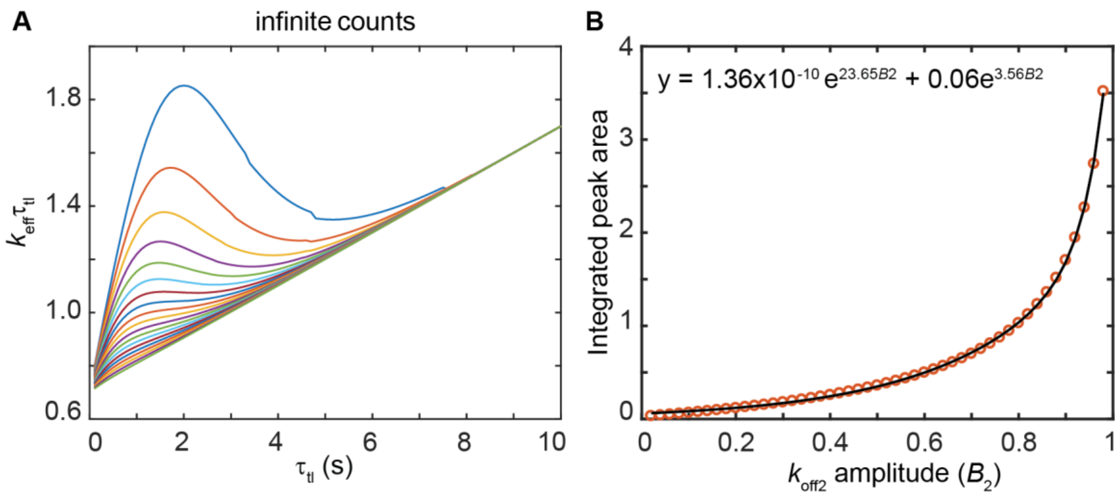


**Figure S5.** Bi-exponential distributions with an intermediate rate ($k_{off1}$ = 0.1 s$^{-1}$) and a fast rate ($k_{off2}$ = 1 s$^{-1}$) with infinite counts. (A) Representative $k_{eff}\tau_{tl}$ plots at 20 amplitudes of $k_{off2}$. From top to bottom, the amplitude reduces from 95% to 5%. (B) Integrated peak areas as a function of $k_{off2}$ amplitudes (open circles). Line is the exponential fit to data points ($R^2$: 0.9996). The peak area is calculated as the difference between areas under the $k_{eff}\tau_{tl}$ plots and the area under the line $y = 0.7 + 0.1\tau_{tl}$.

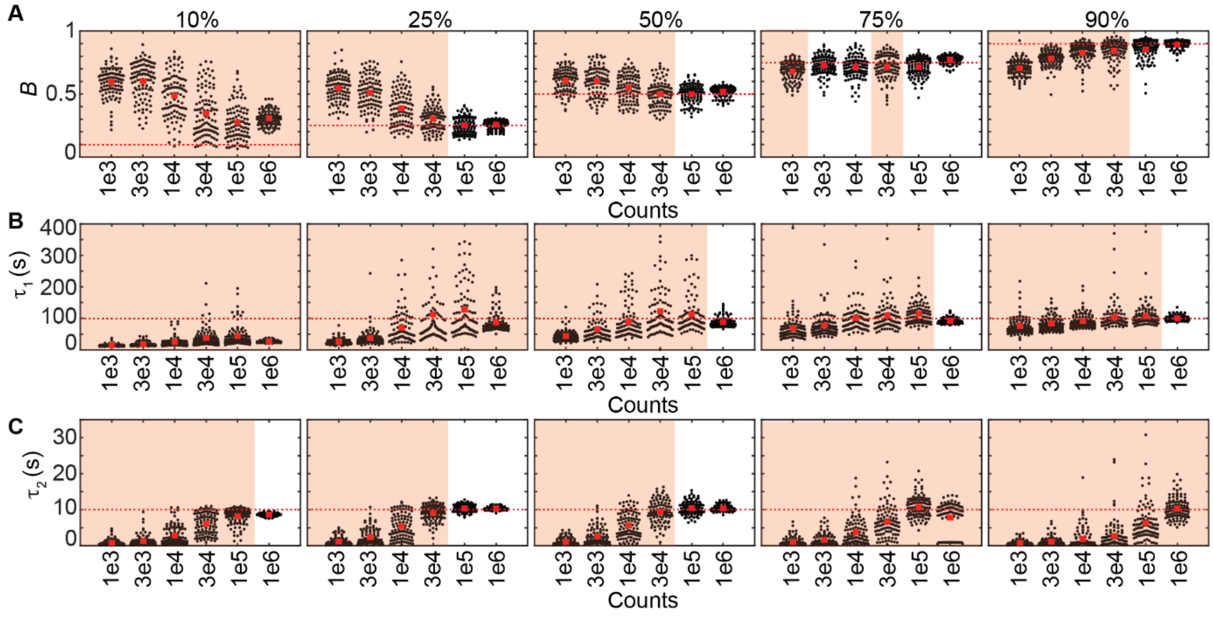


**Figure S6.** Determination of time constants and amplitudes from bi-exponential distributions with a slow rate ($k_{off1}$ = 0.01 $s^{-1}$) and an intermediate rate ($k_{off2}$ = 0.1 $s^{-1}$). (A-C) Scatter plots show distributions of *B*, $\tau_1$ and $\tau_2$ obtained from fitting of 100 simulated distributions to bi-exponential model. Each panel corresponds to a pre-set amplitude of *B*, which increases from 10%, 25%, 50%, 75% to 90% from left to right. In each panel, *n* increases from $10^3$ (1e3) to $10^6$ (1e6). Dashed lines and red bars represent the true values and the average respectively. Orange shades represent distributions where $\sigma_B$ is larger than 0.1 or $\sigma_\tau/\tau$ is larger than 20%. To enhance visibility, outliers (less than 5% when present) were omitted from scatter plots.

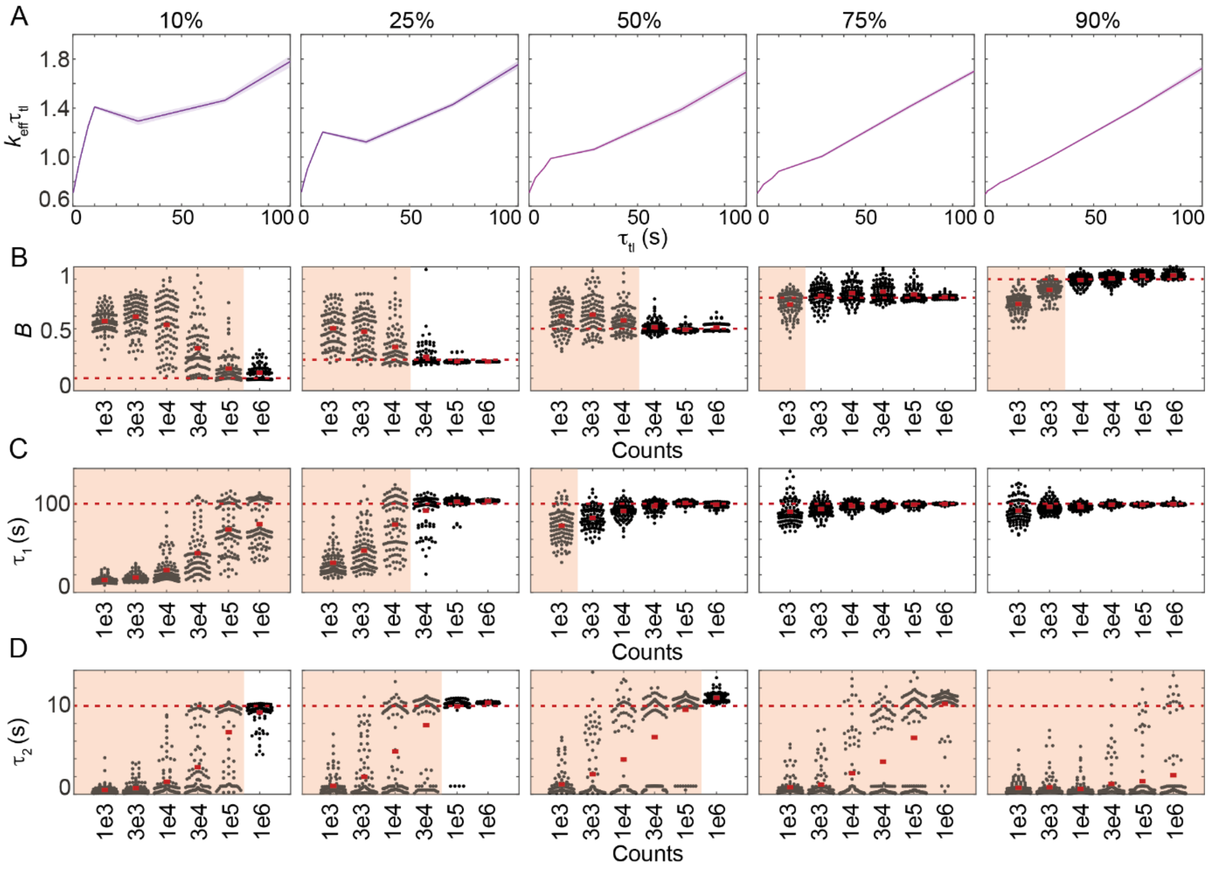


**Figure S7.** Determination of time constants and amplitudes from bi-exponential distributions with a slow rate ($k_{off1}$) and an intermediate rate ($k_{off2} = 10k_{off1}$), simulated using the 100-s $\tau_{tl}$ set. (A) $k_{eff}\tau_{tl}$ plots of bi-exponential distributions with $k_b\tau_{int}$ of 0.7, $k_{off1}$ and $k_{off2}$ of 0.01 and 0.1 $s^{-1}$ respectively, with $10^5$ observations. The amplitude of $k_{off1}$ (*B*, shown on top) increases from left to right (10% to 90%). Shaded error bands are standard deviations from ten bootstrapped samples. (B-D) Scatter plots show distributions of *B*, $\tau_1$ and $\tau_2$ obtained from fitting of 100 simulated distributions to bi-exponential model. Each panel corresponds to a pre-set amplitude of *B*, which increases from 10%, 25%, 50%, 75% to 90% from left to right. In each panel, *n* increases from $10^3$ (1e3) to $10^6$ (1e6). Dashed lines and red bars represent the true values and the average respectively. Orange shades represent distributions where $\sigma_B$ is larger than 0.1 or $\sigma_\tau/\tau$ is larger than 20%. To enhance visibility, outliers (less than 5% when present) were omitted from scatter plots.

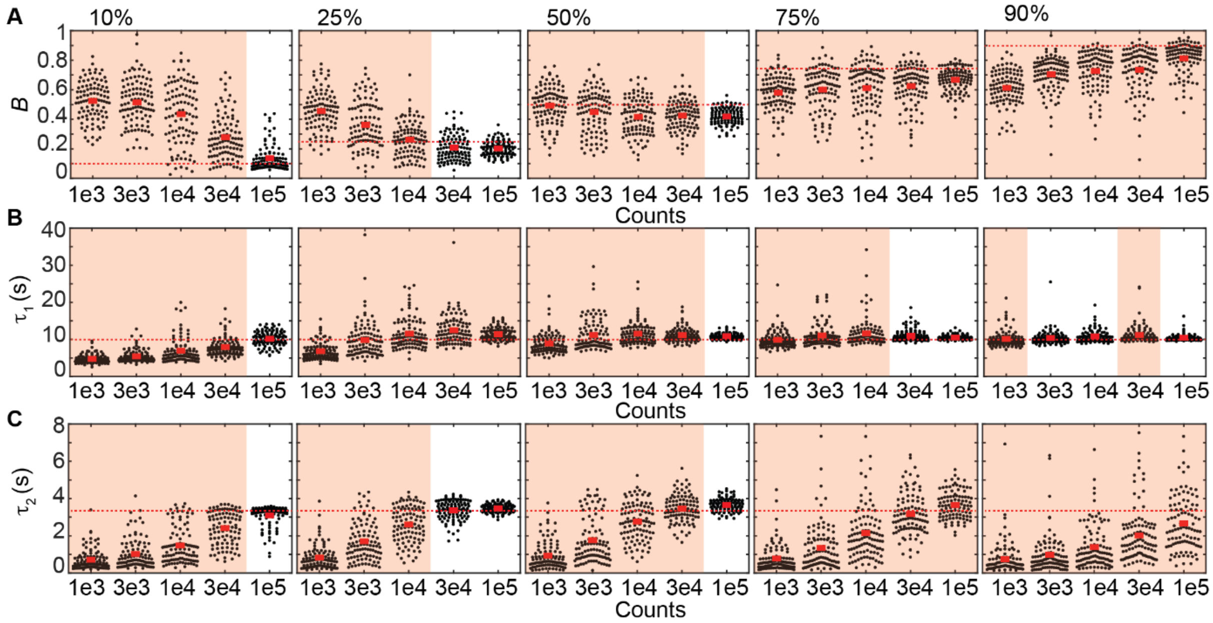


**Figure S8.** Determination of binding lifetimes and amplitudes from bi-exponential distributions with closely spaced rates ($k_{off2}$ = 3$k_{off1}$). (A-C) Scatter plots show distributions of *B*, $\tau_1$ and $\tau_2$ obtained from fitting of 100 simulated distributions for each *n* value. Each panel corresponds to a pre-set *B*, which increases from 10%, 25%, 50%, 75% to 90% from left to right. In each panel, *n* increases from $10^3$ (1e3) to $10^5$ (1e5). Dashed lines and red bars represent the true values and the average respectively. Orange shades represent distributions where $\sigma_B$ is larger than 0.1 or $\sigma_\tau/\tau$ is larger than 20%. To enhance visibility, outliers (less than 5% when present) were omitted from scatter plots.

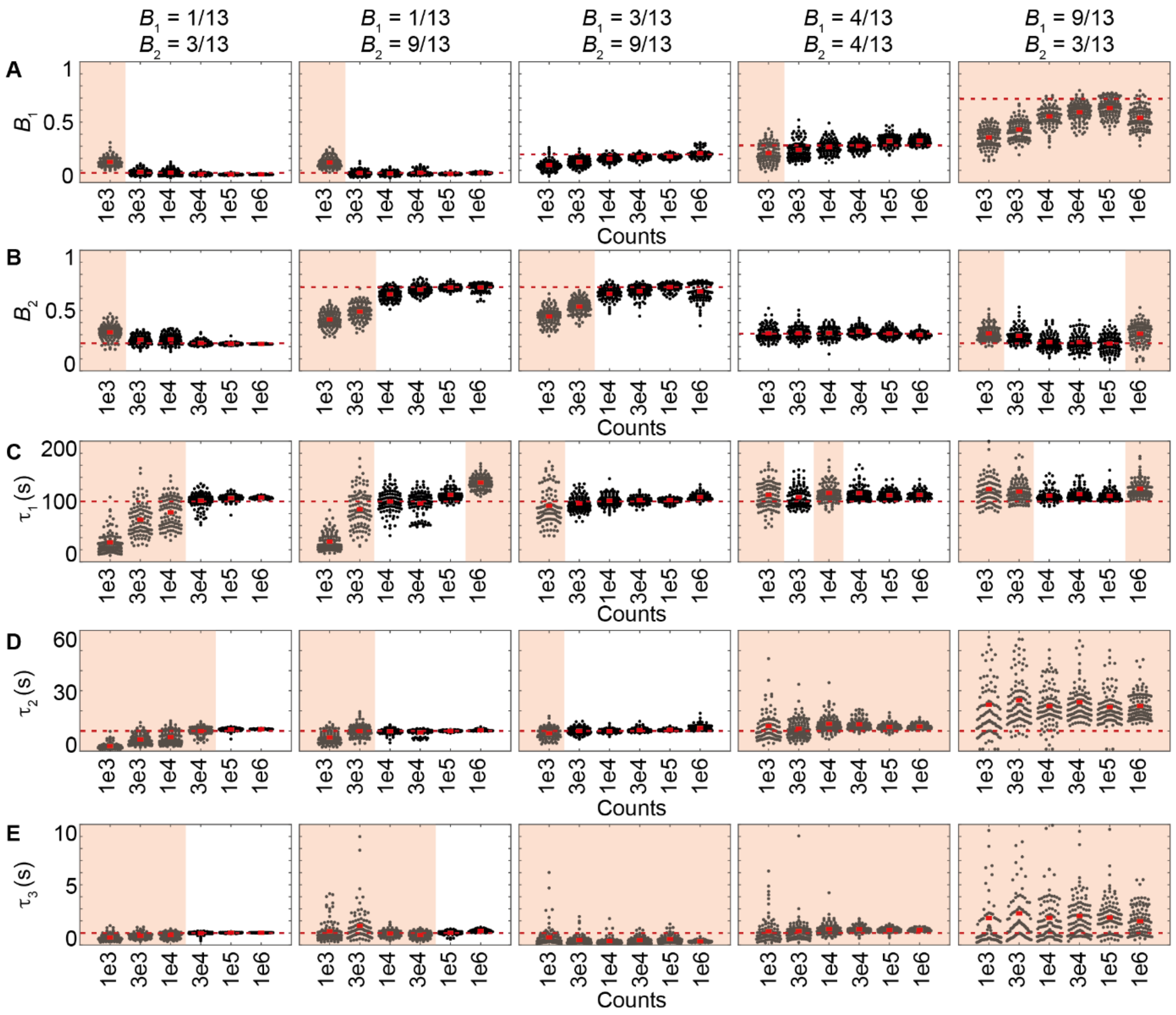


**Figure S9.** Determination of binding lifetimes and amplitudes from tri-exponential distributions with a slow rate ($k_{off1}$), an intermediate rate ($k_{off2} = 10k_{off1}$) and a fast rate ($k_{off3} = 10k_{off2}$), using the 100-s $\tau_{tl}$ set. From left to right, five panels in each row correspond to different amplitudes of each sub-population (displayed on top). (A-E) Scatter plots show distributions of amplitudes ($B_1$ and $B_2$), $\tau_1$, $\tau_2$ and $\tau_3$ obtained using global fitting 100 simulated samples. In each panel, *n* increases from $10^3$ (1e3) to $10^6$ (1e6). Dashed lines and red bars represent the true values and the averages respectively. Orange shades represent distributions where $\sigma_B$ is larger than 0.1 or $\sigma_\tau/\tau$ is larger than 20%. To enhance visibility, outliers (less than 5% when present) were omitted from scatter plots.

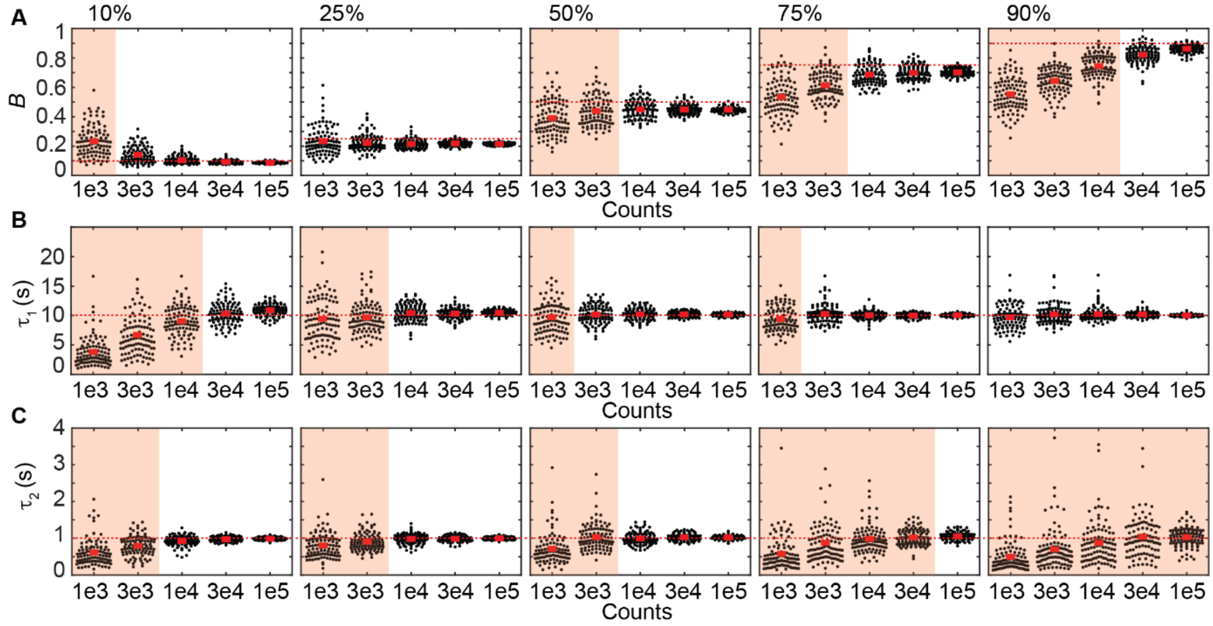


**Figure S10.** Determination of time constants and amplitudes from bi-exponential distributions simulated with the five $\tau_{tl}$ set, and an intermediate rate ($k_{off1}$ = 0.1 $s^{-1}$) and a fast rate ($k_{off2}$ = 1 $s^{-1}$). (A-C) Scatter plots show distributions of $B$, $\tau_1$ and $\tau_2$ obtained from fitting of 100 simulated distributions to bi-exponential model. Each panel corresponds to a pre-set amplitude of $B$, which increases from 10%, 25%, 50%, 75% to 90% from left to right. In each panel, $n$ increases from $10^3$ (1e3) to $10^5$ (1e5). Dashed lines and red bars represent the true values and the average respectively. Orange shades represent distributions where $\sigma_B$ is larger than 0.1 or $\sigma_\tau/\tau$ is larger than 20%. To enhance visibility, outliers (less than 5% when present) were omitted from scatter plots.

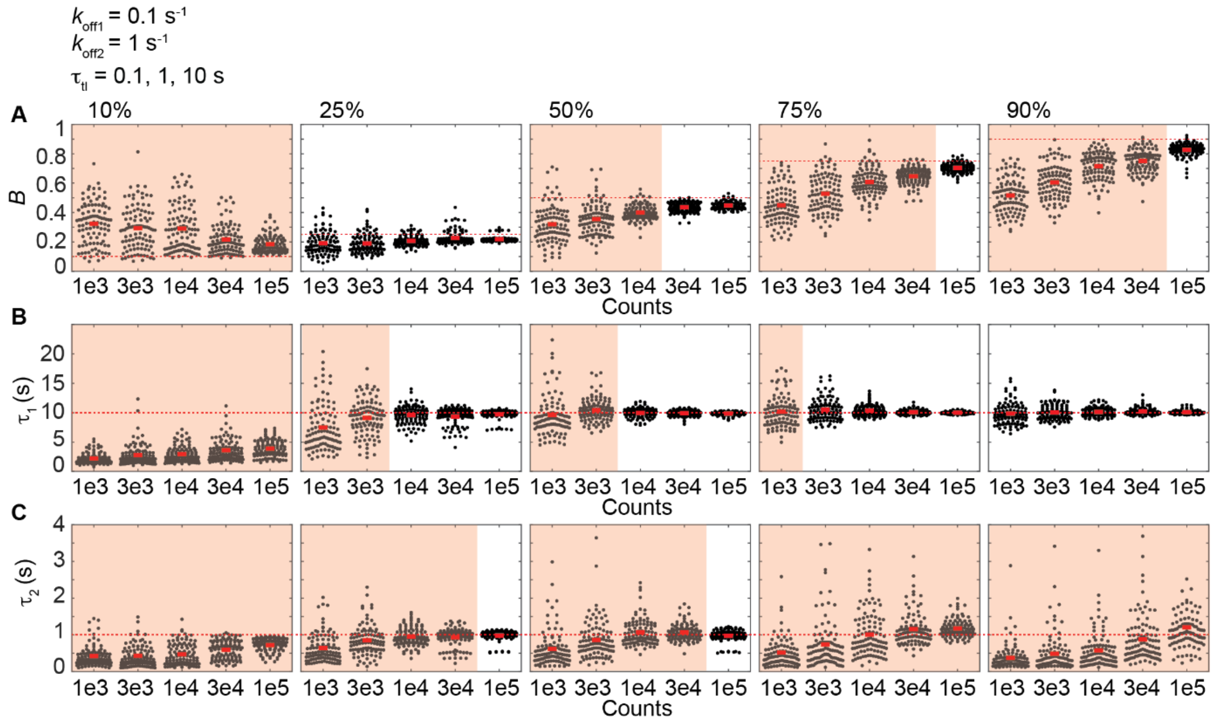


**Figure S11.** Determination of time constants and amplitudes from bi-exponential distributions simulated with the three $\tau_{tl}$ set, and an intermediate rate ($k_{off1}$ = 0.1 s$^{-1}$) and a fast rate ($k_{off2}$ = 1 s$^{-1}$). (A-C) Scatter plots show distributions of *B*, $\tau_1$ and $\tau_2$ obtained from fitting of 100 simulated distributions to bi-exponential model. Each panel corresponds to a pre-set amplitude of *B*, which increases from 10%, 25%, 50%, 75% to 90% from left to right. In each panel, *n* increases from $10^3$ (1e3) to $10^5$ (1e5). Dashed lines and red bars represent the true values and the average respectively. Orange shades represent distributions where $\sigma_B$ is larger than 0.1 or $\sigma_\tau/\tau$ is larger than 20%. To enhance visibility, outliers (less than 5% when present) were omitted from scatter plots.

### Supplementary Notes

1. **Simulation of a set of binding events whose lifetimes follow an exponential distribution with user-defined mean**

```
function [counts, each_molecule] = simulate_res_time(mu,edges,n_count)
%%%%%%%%%%%%%%%%%%%%%%%%%%%%%%%%%%%%%%%%%%%%
%% Inputs:
%%    mu: mean of exponential distribution for a particular τtl
%%    edges: bin edges of histograms
%%    n_count: the number of counts for a particular τtl
%% Outputs:
%%    counts: vector describing CRTD
%%    each_molecule: vector containing all random number corresponding to
%%                   lifetimes of binding events
%%%%%%%%%%%%%%%%%%%%%%%%%%%%%%%%%%%%%%%%%%%%
each_molecule = [];
counts = zeros(10,1);
%% generate a set of random numbers corresponding to lifetimes of binding
%% events until counts in the first bin exceed user-defined counts
while counts(1) < n_count
% single iteration of the exprnd function
    sim = exprnd(mu,round(n_count/2.71),1);
% construct the histogram with edges corresponding to frame times
% N is a vector containing counts in all bins [from the latest iteration]
    [N,~] = histcounts(sim,edges);
    counts = counts + N'; % add counts to the previous iterations
% combine lifetimes of binding events to existing population from previous
% iteration of the exprnd function
    each_molecule = [each_molecule; sim];
end
end % end of the function
```

2. **Simulation of mono-, bi- or tri-exponential distribution across all $\tau_{tl}$**

```
%%%%%%%%%%%%%%%%%%%%%%%%%%%%%%%%%%%%%%%%%%%%
%% Inputs:
%%    ttl:   vector containing the set of time-lapse intervals
%%    kb:    photobleaching rate (unit: s-1)
%%    tint:  camera integration time
%%    koff1: user-defined off rate 1
%%    koff2: user-defined off rate 2
%%    koff3: user-defined off rate 3
%%    B(1):  amplitude of the first kinetic sub-population
%%    B(2):  amplitude of the second kinetic sub-population
%%    n_count_total: user-defined counts for each simulation
%% Outputs:
%%    bin: matrix containing CRTDs for all time-lapse intervals
%%    d.data: contains the simulated population at a particular time-
%%    lapse interval
%%%%%%%%%%%%%%%%%%%%%%%%%%%%%%%%%%%%%%%%%%%%
for i = 1:length(ttl)    % simulate CRTD for each time-lapse interval
     time = ttl(i)*(0:10)';  % determine frame times for binning
     %% define exponential distribution for each sub-population
     keff1 = (kb*tint/ttl(i) + koff1); % effective rate 1
     % mean of the exponential distribution of the first sub-population
     mu1 = 1/keff1;
     keff2 = (kb*tint/ttl(i) + koff2); % effective rate 2
     % mean of the exponential distribution of the second sub-population
     mu2 = 1/keff2;
     keff3 = (kb*tint/ttl(i) + koff3); % effective rate 3
     % mean of the exponential distribution of the third sub-population
     mu3 = 1/keff3;
     %% determine the number of counts for each sub-population based
     %% on the amplitudes B1 and B2
     % counts of the first kinetic sub-population
     n_count1 = round(B(1)*n_count_total);
     % counts of the second kinetic sub-population
     n_count2 = round(B(2)*n_count_total);
     % counts of the third kinetic sub-population
     n_count3 = n_count_total - n_count1 - n_count2;
     % bin1, bin2 and bin3 are vectors containing CRTDs of koff1, koff2 and
     % koff3 sub-population respectively
     % population1, population2 and population3 are vectors containing
     % simulated koff1, koff2 and koff3 sub-population respectively.
     bin2 = zeros(10,1); population2 = [];
     bin3 = zeros(10,1); population3 = [];
     % simulate koff1 sub-population
     [bin1, population1] = simulate_res_time(mu1,time,n_count1);
     % simulate koff2 sub-population
     if n_count2 > 1
          [bin2, population2] = simulate_res_time(mu2,time,n_count2);
     end
     % simulate koff3 sub-population
     if n_count3 > 1
          [bin3, population3] = simulate_res_time(mu3,time,n_count3);
     end
     % combine CRTDs from sub-population CRTDs
            bin(:,i) = bin1 + bin2 + bin3;
     % combine simulated population from simulated sub-populations
     d(i).data = [population1; population2; population3];
end
```

### 3. Global fitting

```
function [p_out] = globalFit(i_model, X, Y, tint)
%%%%%%%%%%%%%%%%%%%%%%%%%%%%%%%%%%%%%%%%%%%%
%% Inputs:
%%    i_model = 1 - mono-exponential model
%%    i_model = 2 - bi-exponential model
%%    i_model = 3 - tri-exponential model
%%    X: matrix containing frame times of all time-lapse intervals
%%          - row: frame times corresponding to one time-lapse interval
%%          - column: increase in frame times
%%    Y: matrix containing simulated CRTDs of all time-lapse intervals
%%    tint: camera integration time
%%    para: initial conditions
%%      - mono-exponential model: [kb, koff1, counts]
%%      - bi-exponential model: [kb, koff1, B1, koff2, counts]
%%      - tri-exponential model: [kb, koff1, B1, koff2, B2, koff3, counts]
%%    lb: lower constraints
%%      - mono-exponential model: [kb, koff1, counts]
%%      - bi-exponential model: [kb, koff1, B1, koff2, counts]
%%      - tri-exponential model: [kb, koff1, B1, koff2, B2, koff3, counts]
%%    ub: upper constraints
%%      - mono-exponential model: [kb, koff1, counts]
%%      - bi-exponential model: [kb, koff1, B1, koff2, counts]
%%      - tri-exponential model: [kb, koff1, B1, koff2, B2, koff3, counts]
%% Outputs:
%%    p_out: vector containing outcomes of global fitting
%%          - p(1): kb
%%          - p(2): koff1
%%          - p(3): B1
%%          - p(4): koff2
%%          - p(5): B2
%%          - p(6): koff3
%%          - p(7): 1 - B1 - B2
%%          - p(8)-p(end): counts at time 0 for all time-lapse intervals
%%%%%%%%%%%%%%%%%%%%%%%%%%%%%%%%%%%%%%%%%%%%
%% Known Parameters
ttl = X(:,1); % vector containing all time-lapse intervals
a_para = Y(:,1); % Initialize the vector for counts at time 0
weights = ones(size(X)); % fitting weights
lower_B = 1/min(a_para(a_para>0)); % the lower bound for the amplitudes
upper_koff = 1/tint;     % the upper bound for off rates
if i_model == 1   % fitting to mono-exponential function
     para = [1,  1, a_para']; % initial conditions: kb, koff1, counts
     lb   = [0,  0, zeros(size(ttl))']; % lower bounds: kb, koff1, counts
     % upper bounds: kb, koff1, counts
     ub   = [Inf,upper_koff, Inf*ones(size(ttl))'];
     % define function to minimize
     f1 = @(p)(   (model(i_model,p,X,tint,ttl)-Y).*weights );
     opts = optimset('Display','off');
     % Global fitting using the lsqnonlin function
     [p] = lsqnonlin(f1,para,lb,ub,opts);
     p_out = [p(1:2),1,zeros(1,4),p(3:end)];
elseif i_model == 2 % fitting to bi-exponential function
     para = [1, 1, 0.5, 2, a_para'];
     lb   = [0, 1e-3, lower_B, 1e-3, zeros(size(ttl))'];
     ub   = [Inf, upper_koff, 1-lower_B, upper_koff, Inf*ones(size(ttl))'];
     % define function to minimize
     f1 = @(p)(   (model(i_model,p,X,tint,ttl)-Y).*weights );
     opts = optimset('Display','off');
```

```
        % Global fitting using the lsqnonlin function
        [p] = lsqnonlin(f1,para,lb,ub,opts);
         % assign the smaller off rate to be koff1
         p_temp = sortrows([p(2) p(3); p(4) (1 - p(3))]);
         p_temp = p_temp';
         p_out  = [p(1), p_temp(:)', zeros(1,2), p(5:end)];
elseif i_model == 3
         para = [1, 0.05, 0.3, 0.5, 0.3, 5, a_para'];
         lb   = [0, 1e-3, lower_B, 1e-3, lower_B, 1e-3, zeros(size(ttl))'];
         ub   = [Inf, upper_koff, 1-lower_B, upper_koff, 1-lower_B,upper_koff,
               Inf*ones(size(ttl))'];
         % define function to minimize
         f1 = @(p)( sum(sum((model(i_model,p,X,tint,ttl)-Y).^2.*weights,2 )));
         opts = optimoptions('fmincon', 'MaxFunctionEvaluations',10000,...
               'MaxIter',3000,'Algorithm','interior-point','StepTolerance',
               1.0000e-9);
         b = 1-2*lower_B;
         A = [0,0,1,0,1,0,zeros(1,size(a_para,1))];
         % Global fitting using the fmincon function
         [p] = fmincon(f1,para,A,b,[],[],lb,ub,[],opts);
         % assign the smallest off rate to be koff1 and the second smallest to
         % be koff2
         p_temp = sortrows([p(2) p(3); p(4) p(5); p(6) (1-p(3)-p(5))]);
         p_temp = p_temp';
         p_out = [p(1),p_temp(:)',p(7:end)];
end
end % end of function
```

#### 4. Define fitting models

```
function f = model(i_model,para,X,tint,ttl)
%%%%%%%%%%%%%%%%%%%%%%%%%%%%%%%%%%%%%%%%%%%
%% Inputs:
%%     i_model = 1 – mono-exponential model
%%     i_model = 2 – bi-exponential model
%%     i_model = 3 – tri-exponential model
%%     para: global parameters
%%     X:     frame times
%%     tint: camera integration times
%%     ttl: time-lapse time
%%%%%%%%%%%%%%%%%%%%%%%%%%%%%%%%%%%%%%%%%%%
% ampl: vector containing counts for all time-lapse intervals
p = tint./ttl; p = p(:);
if i_model == 1
    kb = para(1);
    koff1 = para(2);
    ampl = para(3:end);
    % mono-exponential model
    f = (ampl'*ones(1,size(X,2))).*
        (exp(-((kb.*p + koff1)*ones(1,size(X,2))).*X));
elseif i_model == 2 % bi-exponential model
    kb = para(1);
    koff1 = para(2);
    B1 = para(3);
    koff2 = para(4);
    ampl = para(5:end);
    % bi-exponential model
    f = (ampl'*ones(1,size(X,2))).*(B1.*exp(-((kb.*p + koff1)*
      ones(1,size(X,2))).*X)+(1-B1).*exp(-(kb.*p + koff2)*
      ones(1,size(X,2)).*X));
```

```
elseif i_model == 3
    kb = para(1);
    koff1 = para(2);     B1 = para(3);
    koff2 = para(4);     B2 = para(5);
    koff3 = para(6);
    ampl = para(7:end);
    % tri-exponential model
    f = (ampl'*ones(1,size(X,2))).*
        (B1.*exp(-((kb.*p + koff1) * ones(1,size(X,2))).*X)
        + B2.* exp( -(kb.*p + koff2)*ones(1,size(X,2)).*X )+
        (1-B1-B2).* exp( -(kb.*p + koff3)*ones(1,size(X,2)).*X ));
end
end % end of function
```
